# Ribosome exit tunnel electrostatics

**DOI:** 10.1101/2020.10.20.346684

**Authors:** Marc Joiret, Francesca Rapino, Pierre Close, Liesbet Geris

## Abstract

The impact of the ribosome exit tunnel electrostatics on the protein elongation rate or on the forces acting upon the nascent polypeptide chain are currently not fully elucidated. In the past, researchers have measured the electrostatic potential inside the ribosome polypeptide exit tunnel at a limited number of spatial points, at least in prokaryotes. Here, we present a basic electrostatic model of the exit tunnel of the ribosome, providing a quantitative physical description of the tunnel interaction with the nascent proteins at all centro-axial points inside the tunnel. We show how the tunnel geometry causes a positive potential difference between the tunnel exit and entry points which impedes positively charged amino acid residues from progressing through the tunnel, affecting the elongation rate in a range of minus 40% to plus 85% when compared to the average elongation rate. The time spent by the ribosome to decode the genetic encrypted message is constrained accordingly. We quantitatively derived, at single residue resolution, the axial forces acting on the nascent peptide from its particular sequence embedded in the tunnel. The model sheds light on how the experimental data point measurements of the potential are linked to the local structural chemistry of the inner wall and the shape and size of the tunnel. The model consistently connects experimental observations coming from different fields in molecular biology, structural and physical chemistry, biomechanics, synthetic and multi-omics biology. Our model should be a valuable tool to gain insight into protein synthesis dynamics, translational control and into the role of the ribosome’s mechanochemistry in the co-translational protein folding.

## I. Introduction

Ribosomes are the cells’ manufacturing tools building up proteins. They decode the 61 sense codons from a primary message encrypted in a messenger RNA (mRNA) single molecule, translate it with the help of a set of fewer than 61 transfer RNAs (tRNAs) into 20 amino acids to be sequentially polymerized in a nascent polypeptide that will eventually fold into its final structure. At each elongation cycle, the ribosome incorporates a new amino acid into the nascent protein and translocates to the next codon – shifting along the single stranded mRNA by three nucleotides (triplet). Ribosomes have three binding sites for tRNAs: the aminoacyl (A), the peptidyl (P), and exit (E) sites, each located between the small and the large subunit of the ribosome. The elongation cycle starts with recognition, accommodation by induced fit and proofreading of an aminoacylated tRNA on the A site of the ribosome if the cognate anticodon pairs the codon being read on the mRNA [1, 2]. Elongation proceeds with the binding of the carboxyl terminal end of the peptide acylated to the previous tRNA at the P site to the amino moiety of the amino acid acylated on the tRNA at the A site. The formation of the new peptide bond between the nascent chain and the new amino acid is catalyzed at the peptidyl transferase center (PTC), Fig. 1, by a ribozyme belonging to the large subunit of the ribosome [3]. Two energy rich guanosine triphosphate molecules (GTP) are used and two elongation factors with GTPase activity assist the ribosome during each elongation cycle. For more than five decades, attempts to model protein synthesis and mRNA translation from first principles have been pursued extensively [4–7]. Although the average codon translation rate is rather constant transcriptome wide, estimated at 5.6 amino acid residues per second in eukaryotes, codon translation rates have been shown to vary up to 100-fold across a single transcript [8, 9]. Many factors influence translation speeds across a single transcript (mRNA), including differences in cognate, near-cognate and non-cognate tRNA relative abundance, nascent-chain charged residues inside the ribosome exit tunnel, mRNA secondary structure, proline residues at either A or P site of the ribosome, steric hindrance between contiguous ribosomes translating the same mRNA molecule, and the finite resource of the ribosome pool available in the cell [10–22]. The individual contributions of each of the previous factors to the rate of the translation are difficult to assess quantitatively and separately. Sometimes, depending on the local sequence in the mRNA encrypted message, all these factors interfere and may either antagonize each other or, on the contrary, add up to increase or decrease the rate of translation significantly [12–14, 20, 21, 23]. This hampers our understanding of the dynamics of protein synthesis and specifically of the elongation rate.

**FIG. 1:**
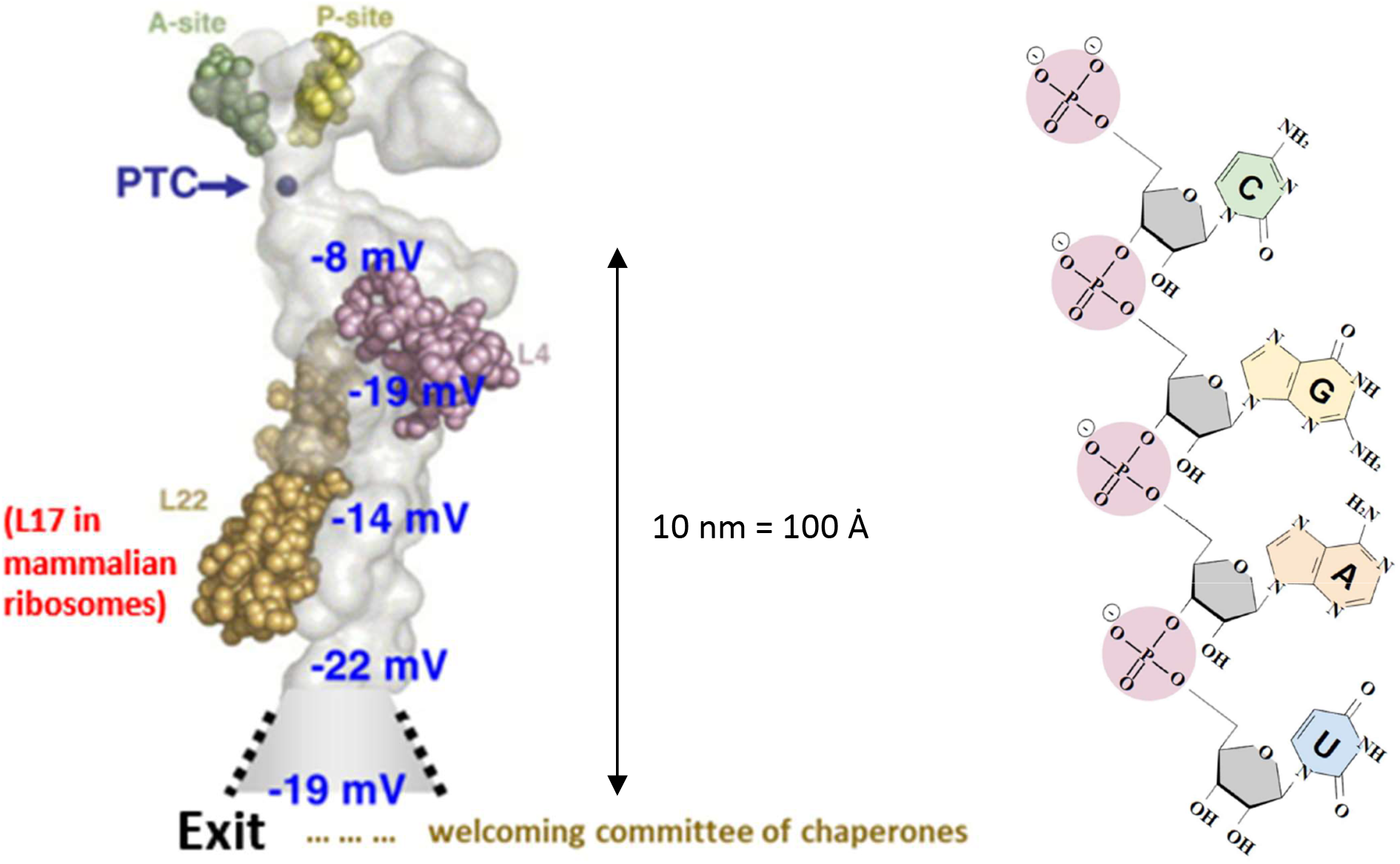
Left panel: Ribosomal exit tunnel structure. The light grey shape is made up of rRNAs. The peptidyl transfer center (PTC) is where a new amino acid residue is bound to the nascent peptide. The figure is taken with permission from Lu *et al* [29]. Right panel: RNA molecular structure showing the ribose-phosphate alternating units. The minus signs represent partial formal negative charges.

Although this has been disputed and it remains a debated question [24–26], some studies have argued that the charged residues are the major determinants of ribosomal velocity [27]. The nascent protein gets out of the ribosome through a tight tunnel approximately 8.5-10 nm long and 1-2 nm wide [28]. The inner wall of the ribosomal exit tunnel is lined with fixed negative charges causing a local negative electrostatic potential inside the tunnel as shown in Fig.1 [29]. Among the 20 amino acids, two of them are positively charged in physiological conditions, namely arginine and lysine [30]. A third one, histidine, is only weakly positively charged. When the ribosome incorporates a local increased number of such positively charged amino acid residues in the nascent protein, a local variation in the elongation rate is often reported. This is also true for the negatively charged amino acid residues, i.e. glutamate and aspartate. Ribosome profiling experiments results (Ribo-Seq) are difficult to interpret and to reconcile with RNA sequencing (RNA-Seq) profiles and proteome expression results in any given biological condition [20]. In vitro laser optical tweezers experiments [31–34], with high resolution dual traps, involving ribosome specifically, are now being conducted to probe the forces acting upon the mRNA at each translocation step or upon the nascent protein emerging from the exit tunnel [32].

The research community would benefit from highly predictive and quantitatively accurate computational models of translation dynamics and specifically of elongation rates. A fully realistic model of the electrostatics inside the ribosome exit tunnel is lacking, despite experimental point measurements of the electrostatic scalar potential in the ribosomal exit tunnel being available, at least in prokaryotes [29]. Stochastic models for protein synthesis have been developed for more than fifty years [4–7, 14, 15]. The extended totally asymmetric simple exclusion process (TASEP) is a widely used stochastic model family dedicated to dynamically simulate the translation rate of a set of transcripts in various conditions [7, 35–37]. In the parametrization of TASEP models, most researchers impose an empirical penalty factor to account for the influence of the electrostatic molecular interaction of the ribosome exit tunnel with newly incorporated charged amino residues at the peptidyl transferase center (PTC). For example, a fixed 20% decrease in the translation rate is imposed for those codons that are within five positions downstream of a codon encoding positively charged amino acid residues (lysine, arginine or histidine)[14]. The negatively charged residues (aspartate and glutamate) are ignored in most studies. This approach is considered inconsistent or too naive if more accurate predictions are expected from TASEP models and to be compared to specific ribosome profiling experimental data [17] or to real-time specific single RNA molecule translation dynamics experiments in vivo [9] or in vitro [31, 32].

In this study, we focus on one of these specific factors which affects the local speed of elongation during protein synthesis, namely the electrostatic interaction between the charged amino acid residues embedded in the nascent polypeptide chain and the ribosome exit tunnel. To model this electrostatic interaction, we developed a full analytical expression of the electrostatic potential inside the tunnel, starting from two very basic and idealized theoretical geometries for the tunnel. The model is used to explore the physical consequences of a possible dynamically variable geometry of the tunnel from a theoretical perspective. The model is used to quantitatively estimate the profile of the axial forces and requires knowledge of the primary sequence of a significant length of the nascent polypeptide chain or its encrypted mRNA to compute the local axial forces acting at the PTC center during elongation. An algorithm is proposed to compute the axial forces acting locally at the PTC and due to a spatially extended electrostatic interaction inside the tunnel. The model is used to conduct comparative analyses of the axial force profiles for different synthetic or real protein sequences. Knowing the axial forces quantitatively allows to estimate the mechanical work and the biochemical energy required at each elongation step to overcome the electrostatic potential barrier inside the ribosome exit tunnel. These estimations are compared to the energy sources and uptakes involved in the mechanochemistry of the ribosome at each elongation cycle. The ribosome exit tunnel electrostatic model we describe can stand as a building block for computational tools that should be beneficial for the analysis of different experimental techniques like the probing of force by laser optical tweezers, the study of conformational changes with fluorescence resonance energy transfer at the ribosome subunits, and for bioinformatic processing of multi-omics data.

## II. Idealized Electrostatic Models of the Ribosomal Exit Tunnel

The local negative electrostatic potential inside the ribosome exit tunnel, shown in Fig.1 (left panel), from which the nascent proteins emerge, originates from the ribosome composition. Ribosomes are composed of two subunits: 50S and 30S in prokaryotes, 60S and 40S in eukaryotes, identified by their sedimentation coefficients, measured in Svedberg unit *S*; the whole prokaryotic and eukaryotic ribosomes are 70S and 80S, respectively. The ribosome exit tunnel is found in the larger (50S or 60S) of the two subunits. Each of the subunits entails proteins and ribosomal ribonucleic acids (rRNAs). The essential feature of interest of rRNAs, shown in Fig. 1 (right panel), is that, like all RNAs, they are single stranded polymerized molecules with a backbone made up of alternating ribose sugars and phosphate groups all esterified alternatively together. In this long strand, each phosphate group harbors a partial negative charge. The inner wall of the ribosome exit tunnel is mainly lined up with rRNAs (more than 80% w/w in eukaryotic ribosome exit tunnels), though in some locations specific proteins are also present.

### A. Hollow straight cylinder model

In a first simplified approach, the ribosome exit tunnel is considered a hollow straight cylinder (Fig. 2 left panel). The wall material is not of the conductor type with mobile free charges but is rather a dielectric material harboring fixed partial charges – the fixed phosphate moieties lining the inner wall. As a first reasonable assumption, the fixed charges are supposed to be uniformly distributed on the surface of the inner wall. The size of the hollow cylinder closest to the shape of the ribosome exit tunnel documented in the literature would be 85 − 100 Å (8.5 − 10 nm) in length and 10 − 20 Å (1 − 2 nm) in diameter [28, 38]. The precise length for the ribosomal exit tunnel as measured by cryo-electron microscopy is 9.2 nm on average in prokaryotes and 8.3 nm on average in eukaryotes [38]. The in-vivo lengths are believed to be a bit larger due to thermal dilatation at the higher temperatures prevailing in living organisms as compared to the cryogenic conditions.

**FIG. 2:**
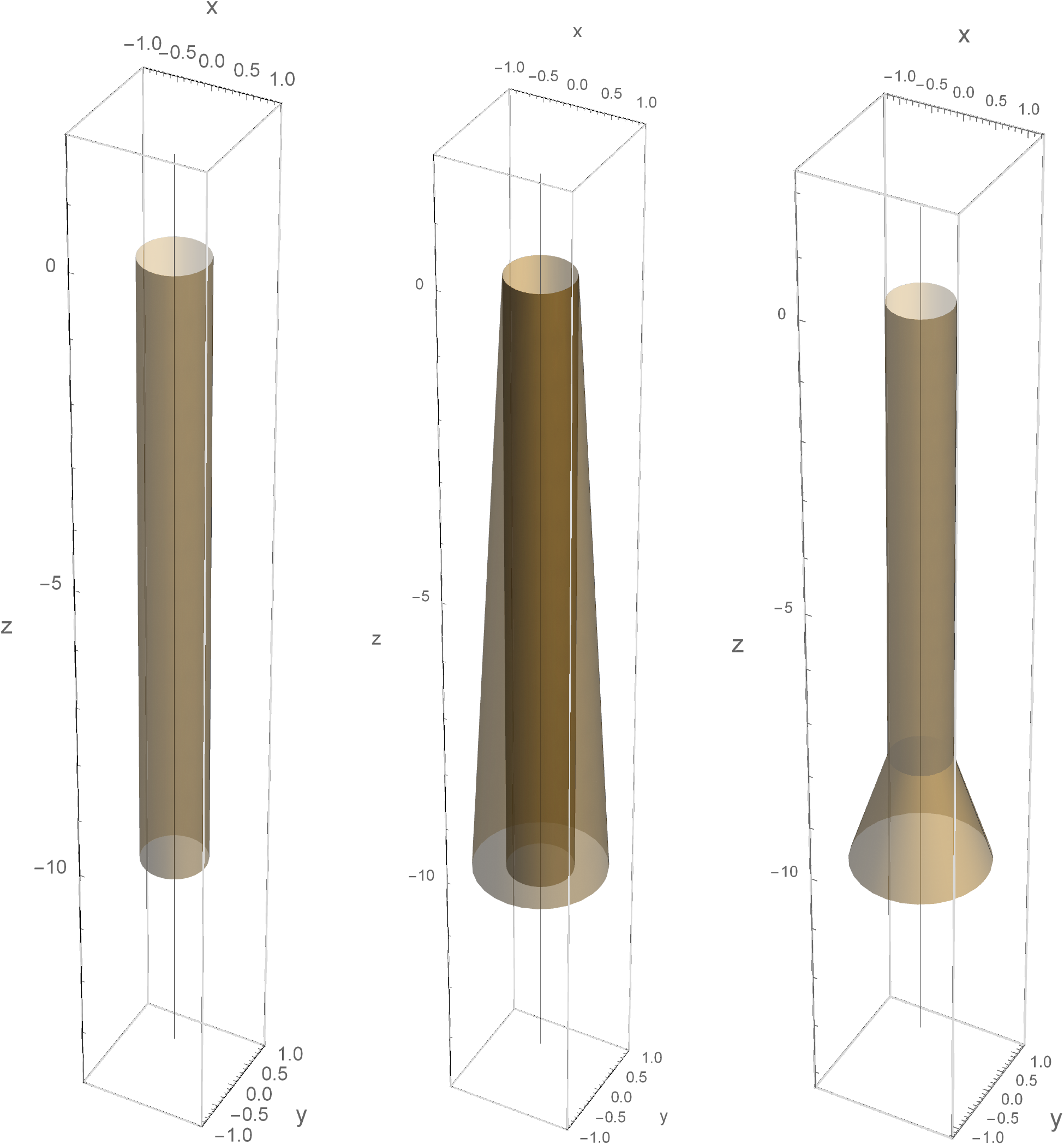
Left panel: hollow cylinder of length L = 10 nm and R = 0.5 nm with uniformly charged inner wall. Center panel: normally truncated cone of length L = 10 nm, R_in_ = 0.5 nm and R_out_ = 1.0 nm with uniformly charged inner wall. The entry point is at *z* = 0 and exit point at *z* = −*L*. Right panel: hollow cylinder of length L_1_ = 8 nm and R = 0.5 nm concatenated to a truncated cone of length L_2_ = 2 nm with a uniformly charged inner wall. The entry point is at *z* = 0 and exit point at *z* = −(*L*_1_ + *L*_2_) = −10 *nm*. Transition between cylinder and truncated cone at 80% of the tunnel total length: *λ* = 0.8.

For a given uniformly distributed charge density *σ* on the inner surface wall of the cylinder, the determination of the electrostatic scalar potential 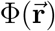 and of the electric field 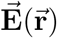, at any spatial point close to or far away from the cylindrical surface, are well stated problems in classical electromagnetism [39]. For the sake of simplicity, we restrict ourselves here on spatial points located on the axis of the hollow cylinder, lying anywhere inside or outside of the tunnel. In this schematic pictorial description, a new amino acid is incorporated into the nascent protein which gets into the tunnel from one side (conventionally from the top of Fig. 2). The nascent oligopeptide is then pushed by the multi-tasking ribosomal enzymatic functions inside the tunnel and out of the tunnel at the other side (bottom side of Fig. 2) of the hollow cylinder. The movement is strictly asymmetric as the nascent protein always enters the tunnel from the same side with the amino terminal end of the protein getting in first and the carboxy terminal end of the protein getting in last. Under this idealized model, the hollow cylinder itself is symmetric and has a uniform charge distribution.

The electrical scalar potential 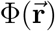 at the observed position 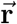 is expressed by:

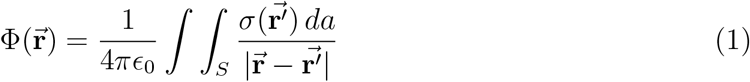

where 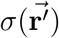 is the surface-charge density (measured in coulombs per square meter) at position 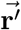 of the source, *da* is the two dimensional surface element at 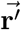 and ϵ_0_ is the permittivity of free space (formula 1.23 in Jackson [39]). We can take advantage of the axial symmetry and restrict to the spatial points on the *z* axis, i.e. for 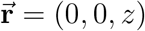. The surface integration is conducted on the support of the source charges. The cylinder’s thin wall is geometrically generated by the *γ*(*u*) curve moving axially along the z-axis from *z* = −*L* to *z* = 0 as drawn in Fig. 2 (left panel) and where *L* and *R* are the length and radius of the hollow cylinder respectively:

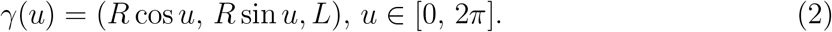

The cylinder’s surface is written as *S* = *ϕ*(*K*) where *K* = {(*u, v*) ∈ [0, 2*π*] × [−1, 0]} and where *ϕ*: ℝ^2^ → ℝ^3^: *ϕ*(*u, v*) = (*R* cos *u, R* sin *u, vL*). *D_u_ϕ* is the first partial derivative of the parametric equation of the surface *ϕ*(*u, v*) with respect to *u*. In the general formula (1), the surface-charge density 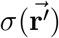 is dependent of the position 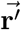 on the support of the source charges. Here, we will take the simple approximation that *σ* can be considered a constant parameter over a surface of a given shape, e.g. over a cylinder or over a cone. This is the surface charge uniform distribution assumption for a given shape.

The electrostatic scalar potential results from the surface integral calculation:

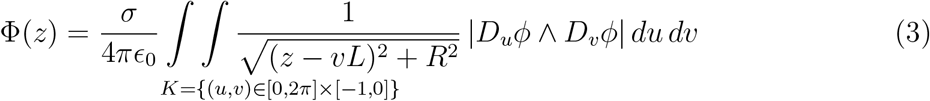

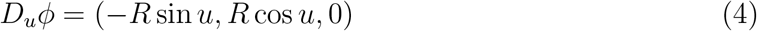

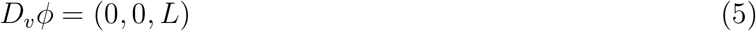

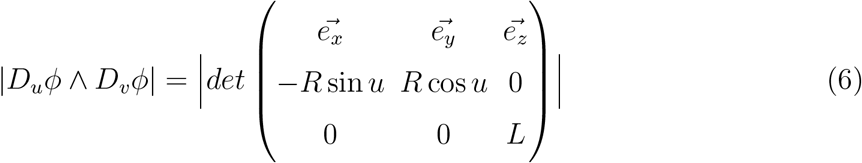

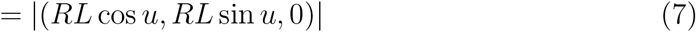

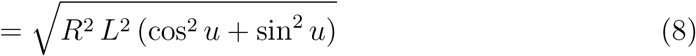

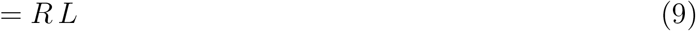

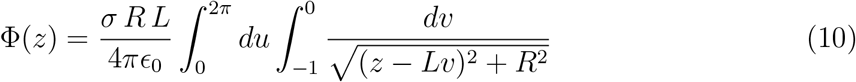

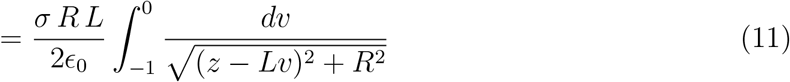

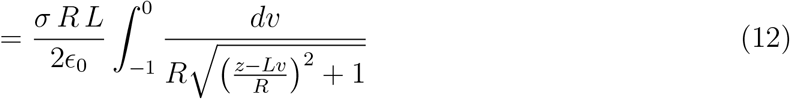

The substitution 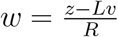 yields 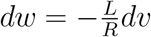 and

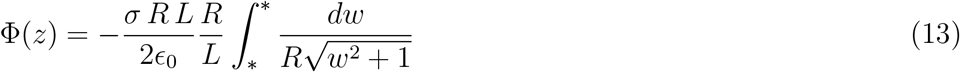

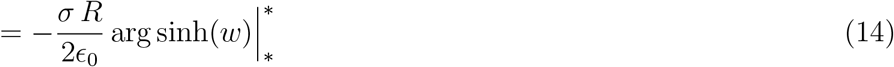

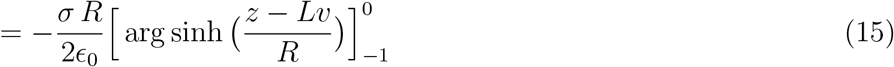

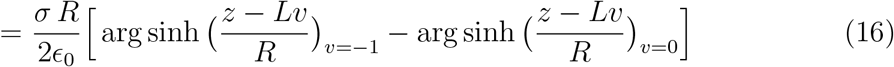

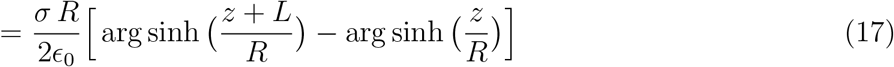

As the arg sinh may be expressed as a logarithm (to prove this, recall that if *x* = sinh *y*, *y* = arg sinh *x* and so 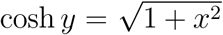 whence, sinh *y* + cosh *y* = *e^y^* and we conclude that 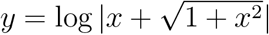, the electrostatic scalar potential finally writes:

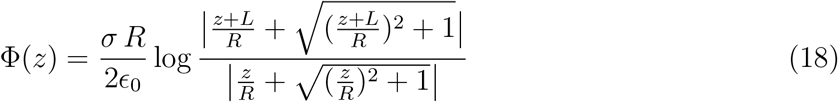

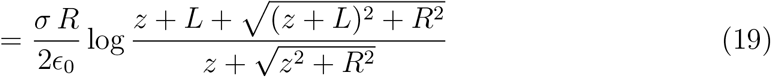

The electric field projected along the cylinder axis can be computed as the opposite of the scalar potential gradient, i.e. by taking the first derivative with respect to *z* directly from formula (17):

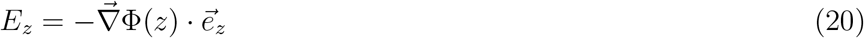

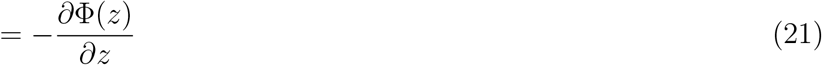

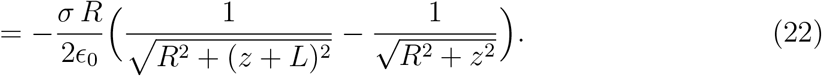

Of course the axial force applied on a test particle is the product of the axial electric field with the charge of the test particle:

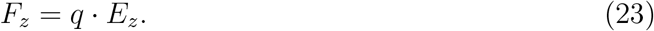

The plots of electrostatic scalar potential Φ(*z*) and of the axial force *F_z_* acting on a unit test charge located on the tunnel axis at any point of coordinate *z* are displayed in Fig. 3, with the medium permittivity prevailing inside the ribosome exit tunnel (see below). A negative force means that the test particle is forced to move towards negative *z* values whereas a positive force means that the test particle is forced to move towards positive *z* values. In these plots, *σ* is adjusted so that the potential fits the range of the experimentally measured values given for instance in Lu *et al.* [29].

**FIG. 3:**
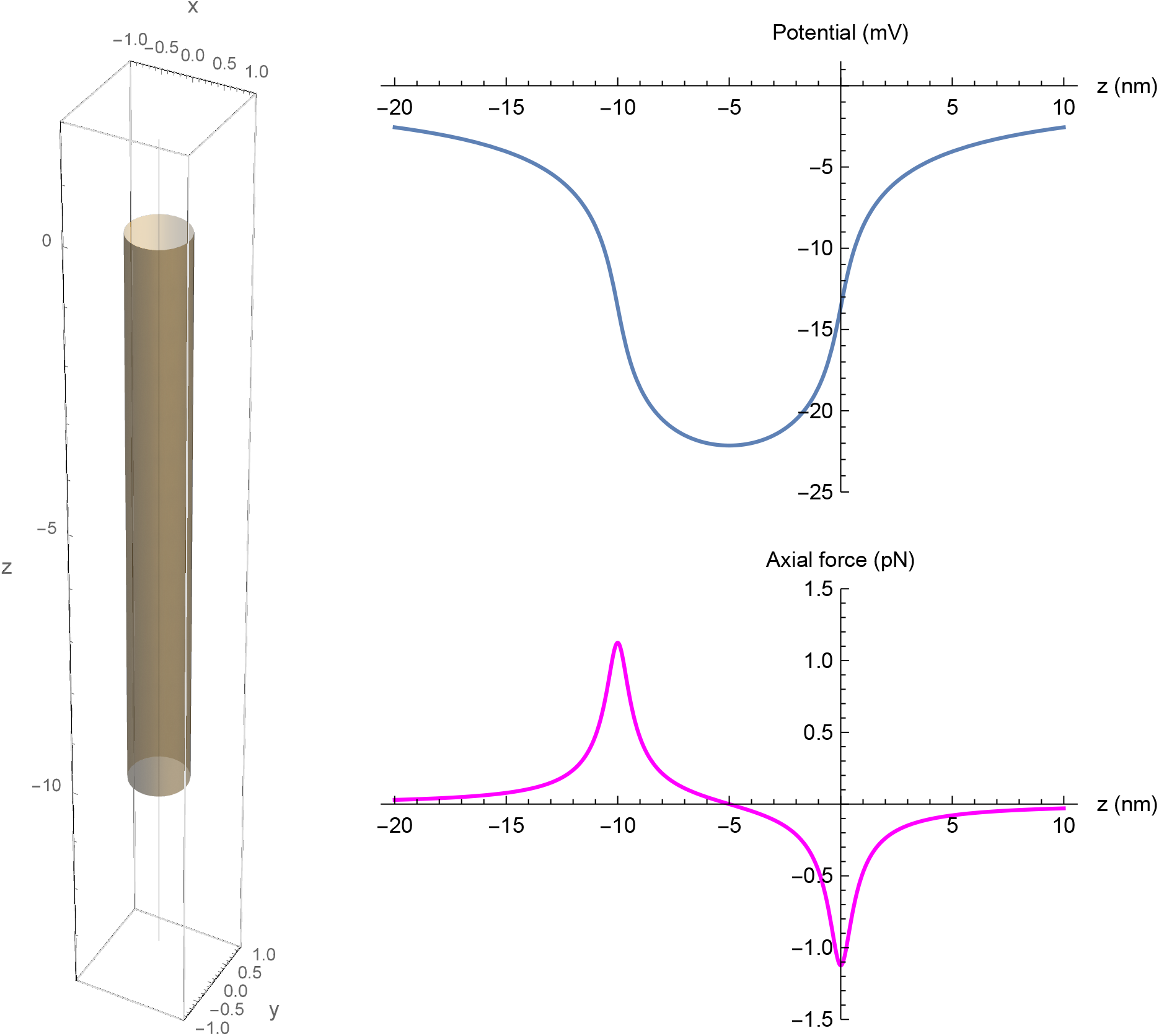
Electrostatic scalar potential on the axis of the ribosomal exit tunnel (upper panel) and axial force (lower panel) as a function of axial *z* position for a positively unit charged test amino acid residue on the tunnel axis. Tunnel idealized as a cylinder.

### B. Normally truncated straight cone model

An alternative approach would depict the tunnel as a hollow cone normally truncated at both ends (Fig. 2 center panel). The section radius at the entry point is still equal to R = 0.5 nm but with a section radius twice that value at the tunnel exit point, and equal to R = 1 nm. With the total axial length kept at L = 10 nm, the half opening angle along the axis is *α* ~ 0.05 radian (2.86 arc degrees) and exactly such that tan *α* = R/L complying with the observation that the diameter at the exit point is around twice the diameter at the entry point of the tunnel. This better reflects the actual geometry of the real ribosomal exit tunnel as reported in the literature [28].

To analytically derive the correct equation for the potential and axial electrical field in such a conical tunnel, the procedure is the same as the one previously conducted for the cylinder, but this time with the support of the uniformly distributed charges defined by a cone surface normally truncated at both ends.

The cone’s surface is written as *S* = *ϕ*(*K*) where *K* = {(*u, v*) ∈ [0, 2*π*] × [−1, 0]} and where 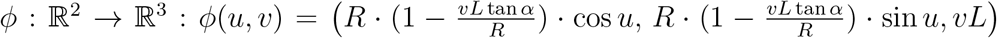. The electrostatic scalar potential results from the surface integral calculation:

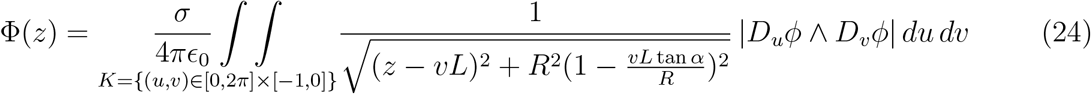

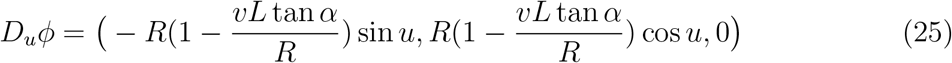

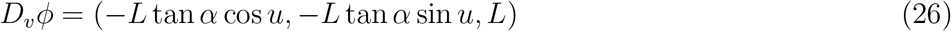

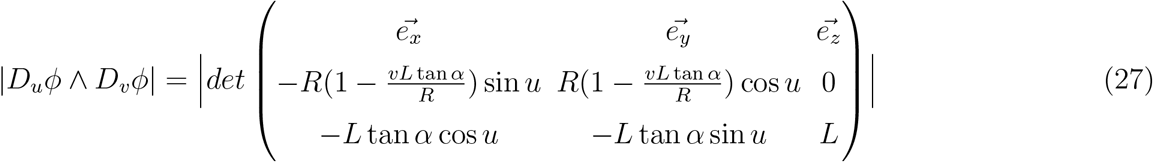

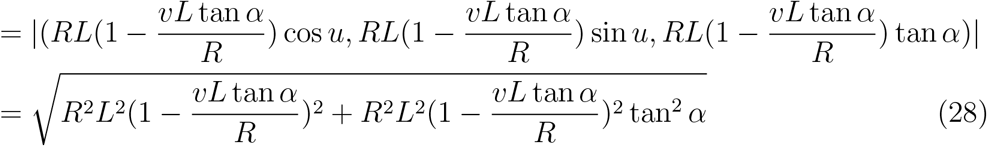

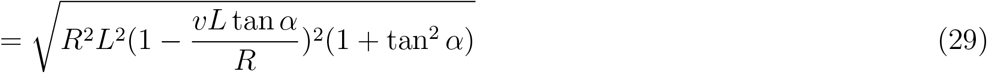

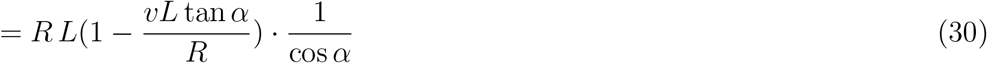

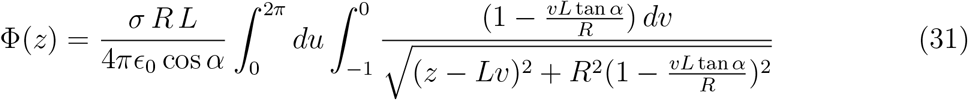

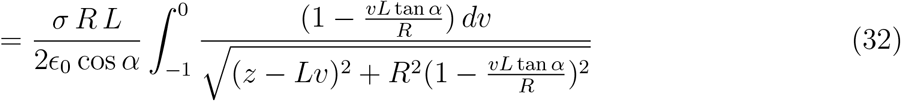

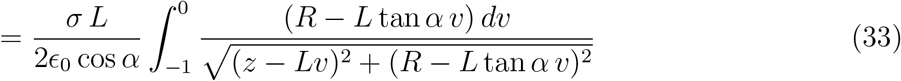

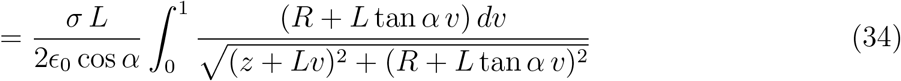

where, in the last line, a dummy integration variable was changed with v′ = −v → dv′ = −dv and the change of sign was cancelled by the integration limits permutation. The complete derivation is given in the supplemental materials.To alleviate the notations, the two following substitutions are adopted:

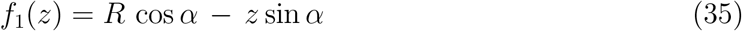

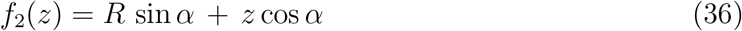

*f*_1_(*z*) is always positive for *z* ≤ 0 (and even for *z < R/* tan *α*, i.e. the virtual *z* position of the cone summit), which is the domain we are interested in. The *z* position values are negative in the tunnel and beyond its exit point.

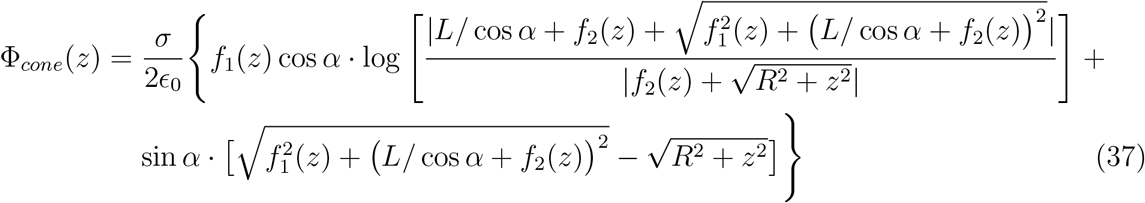

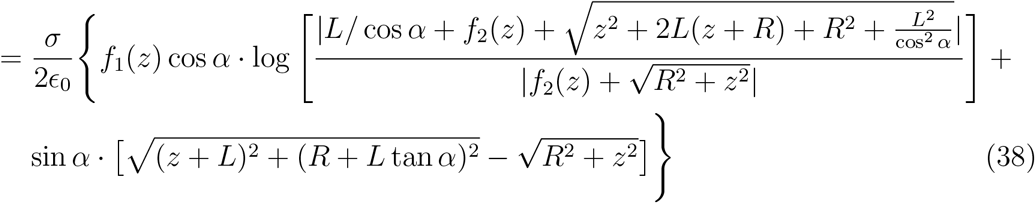

This last equation (38), valid for any conical geometry with entry section of radius *R* and any cone angle *α*, replaces equation (19) of the cylindrical geometry. Note that the electrostatic potential vanishes at *z* = ±∞ as physically expected.

It is also worth noticing that equation (38) for the truncated cone restores, as a special case, equation (19) for the cylinder when *α* = 0, as expected as well.

The electric field projected along the truncated cone axis can be computed as the opposite of the scalar potential gradient, i.e. by taking the first derivative with respect to *z* of equation (38). The full derivation is provided in the supplemental materials and the final result is:

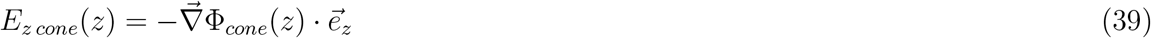

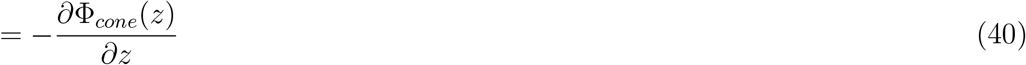

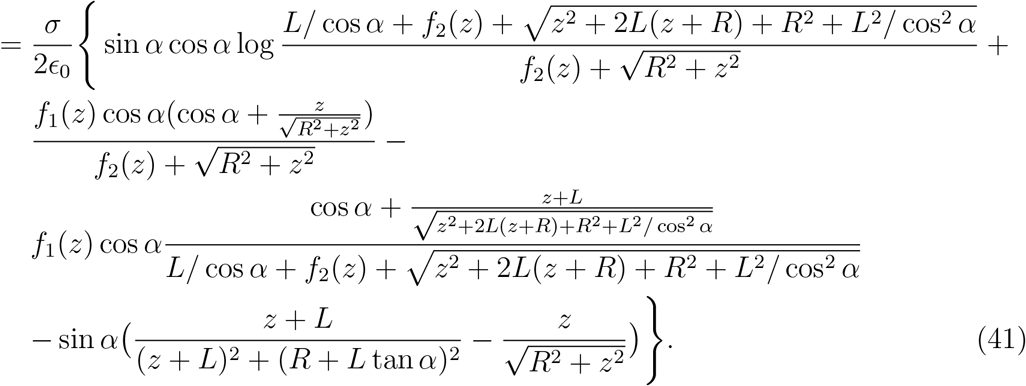

Multiplying eq. (41) by a positive unit test charge yields the axial forces acting on a positive unit test charge. The plot of the axial forces as a function of the position in the tunnel is displayed in Fig. 4 (lower panel) for the truncated cone geometry and compared to the cylinder case.

**FIG. 4:**
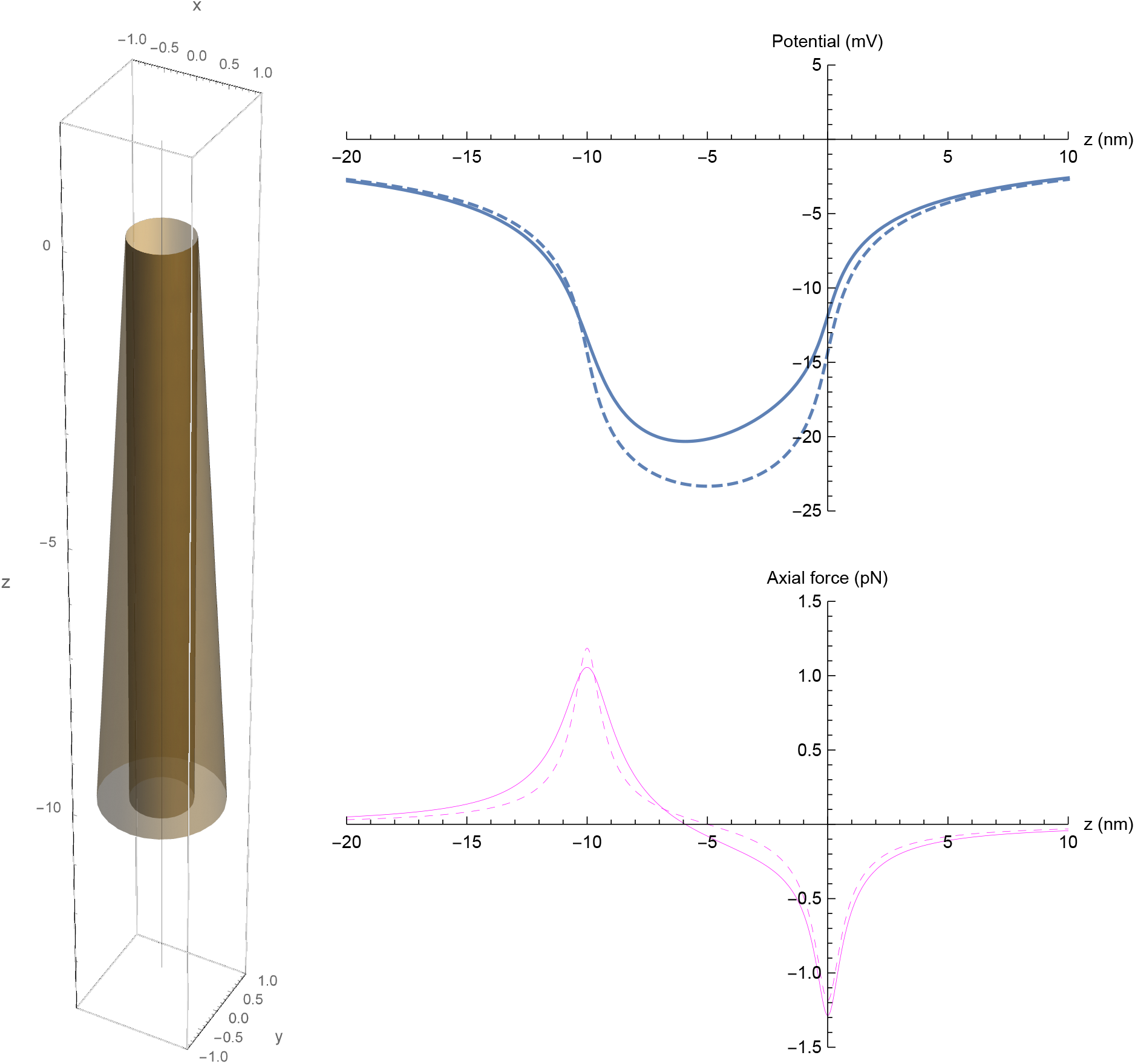
Electrostatic scalar potential and axial force profiles for a positively unit charged test amino acid residue on the axis of the ribosomal exit tunnel. Comparison of the truncated cone (line) and cylinder (dashed line) geometry with exit section radius of the cone twice as large as the cylinder radius. The lateral surface of the cone is 3/2 the lateral surface of the cylinder. *σ*_2_ = *σ*_1_ × 2/3 to keep the same total charges on both surfaces.

Experimental measurements made on ribosome exit tunnels show that the tunnel exit section radius is around 1 nm, i.e. twice the radius of the innermost part of the tunnel. If the ribosome tunnel were of the cone type, the cone opening angle would be around *α* ~ 0.05 radian (2.86 arc degrees).

The consequence on the electrostatic potential profile is of importance because, with this conical geometry, and if the total charges are kept the same for the two surfaces, the electrostatic potential inside the tunnel will necessarily be algebraically higher than the potential profile in the case of the cylinder as displayed in Fig. 4 (upper panel) where the analytical equation for the electrostatic potential for the truncated cone was plotted and compared to the cylinder case.

A simple geometrical calculation shows that if the two surfaces support the same total charges *Q*_1_ = *Q*_2_, then 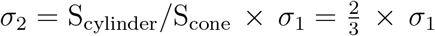, for a geometry where both tunnels have the same radius at the entry point, the same total lengths *L*, but where the cone exit section has a radius twice as large as the cylindrical radius. The surface charge density *σ*_2_ on the lateral truncated cone inner surface would be two third of the surface charge density *σ*_1_ prevailing on the lateral inner surface of the cylinder.

Moreover, the potential profile in the conical geometry is skewed to the left as compared to the potential profile for the cylindrical geometry. An asymmetry in the potential profile appears due to the change in radius along the z-axis of the cone. The minimal value of the potential is shifted to the left. The slope of the cylindrical potential profile is steeper than the conical potential at the tunnel exit point, meaning that the electric field intensity will be a bit weaker in that region for the conical geometry as can be seen in Fig. 4 (lower panel) of the axial forces curves. The axial forces vary more smoothly and are more dispersed in the conical geometry than in the cylindrical geometry.

### C. Normally truncated cone concatenated to a cylinder

The question to know whether or not the tunnel is geometrically exactly more like a cylinder or like a truncated cone is less important than the consequence on the electrostatic potential profile. The salient feature of the real ribosome exit tunnel is that there is indeed a widening in the tunnel section at the exit.

A still better simple geometrical model that fits most of the experimental observations to date is a model combining the cylinder and the normally truncated cone as shown in Fig. 2 (right panel).

The transition from a cylindrical shape to a conical shape results in an electrostatic potential rise along the tunnel axis when moving from the entry point to the exit point. This is a fundamental difference between the two geometries (truncated cone combined to cylinder versus cylinder alone) that has both energetical and biological consequences.

The electrostatic potential resulting from such a configuration results from the superposition of the integrand in equation (3) and the integrand in equation (24) to yield

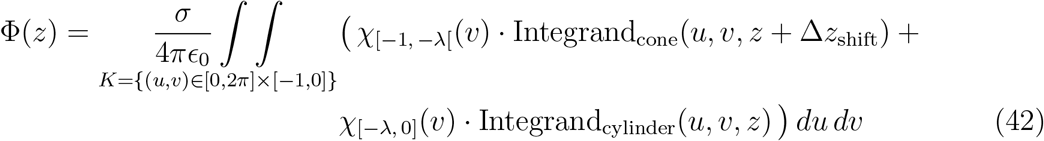

where the characteristic function *χ*_[*a*, *b*]_(*v*), used here for the correct setting of the charged sources distribution, is defined by

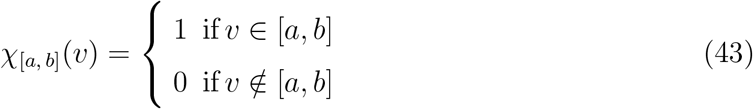

A z-shift was also incorporated in equation (42) to account for the shift in axial position of the truncated cone, as can be noticed by comparing the center and the right panel in Fig. 2. The z-shift must match the *λ* value adopted as the interval limit in both characteristic functions in equation (42), delineating the limit between the start of the truncated cone and the end of the cylinder when moving axially to the left from *z* = 0. For Fig. 2 (right panel), *z*_shift_ = *λ* · *L* = 0.8 *L* (*L* is the tunnel total axial length). The *α* angle for the cone may be a free parameter to be determined. Different surface charge densities can be incorporated as well, using two different values for *σ* in the two integrands, providing an extra degree of freedom to fit the model to the experimental observations. In summary, four parameters can be fitted to the experimental data: *σ*_cylinder_, *σ*_cone_, *λ*, and *α* which are the surface charge density on the cylindrical surface, the surface charge density on the cone frustum, the fraction of the ribosome length occupied by the cylinder, and the cone frustum half opening angle respectively. Each of these parameters can influence the electrostatic potential profile of the tunnel.

The axial electrical field resulting from the combination of the cylindrical geometry for 75% of the tunnel length (0.75 *L*) from its entry point and of the truncated cone geometry for the remaining 25% (0.25 *L*) of the length in the ribosome exit tunnel, with or without an added Lorentzian peak (see below), is the superposition of equations (22) and (41). The parameter settings have to be consistent with the chosen geometry and with the surface charge densities (*σ*_1_ and *σ*_2_). More specifically, for a given *λ*, one would have *L*_1_ = *λ* · *L* and *L*_2_ = (1 − *λ*) · *L*. The surface charge density *σ*_2_ of the truncated cone that we adopted was such that *σ*_2_*/σ*_1_ = *S*_1 lateral_*/S*_2 lateral_ because it best fits the observational data that were gained from the bacteria ribosomes. This condition is consistent with the possibility that the conical part of the tunnel end could result from an elastic deformation of an initially cylindrical shaped tunnel with a uniform surface charge density (conservation of total initial charge before and after this hypothetical elastic deformation of the inner surface turning the cylinder into a truncated cone at the exit side of the tunnel).

The area under the curve of the axial forces profile yields the mechanical energy required for a unit charge to move between two axial points inside the tunnel. Equivalently, the required mechanical energy can easily be computed by multiplying the unit charge with the electrostatic potential difference between the tunnel exit point and the tunnel entry point.

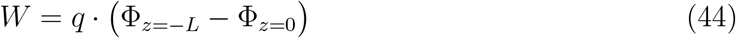

where *W* is the mechanical work required for a unit test charge *q* to move across an electrostatic potential difference (in Volt units) from the tunnel entry point (*z* = 0) to the tunnel exit point (*z* = −*L*). The result is expressed in J/mol units by multiplying equation (44) with the Avogadro number.

The geometrical asymmetry induced by the widening in the tunnel radius at the exit of the tunnel is important because it introduces a permanent difference in the electrostatic potential between the exit and the entry points of the tunnel as shown in Fig. 5 (upper panel).

**FIG. 5:**
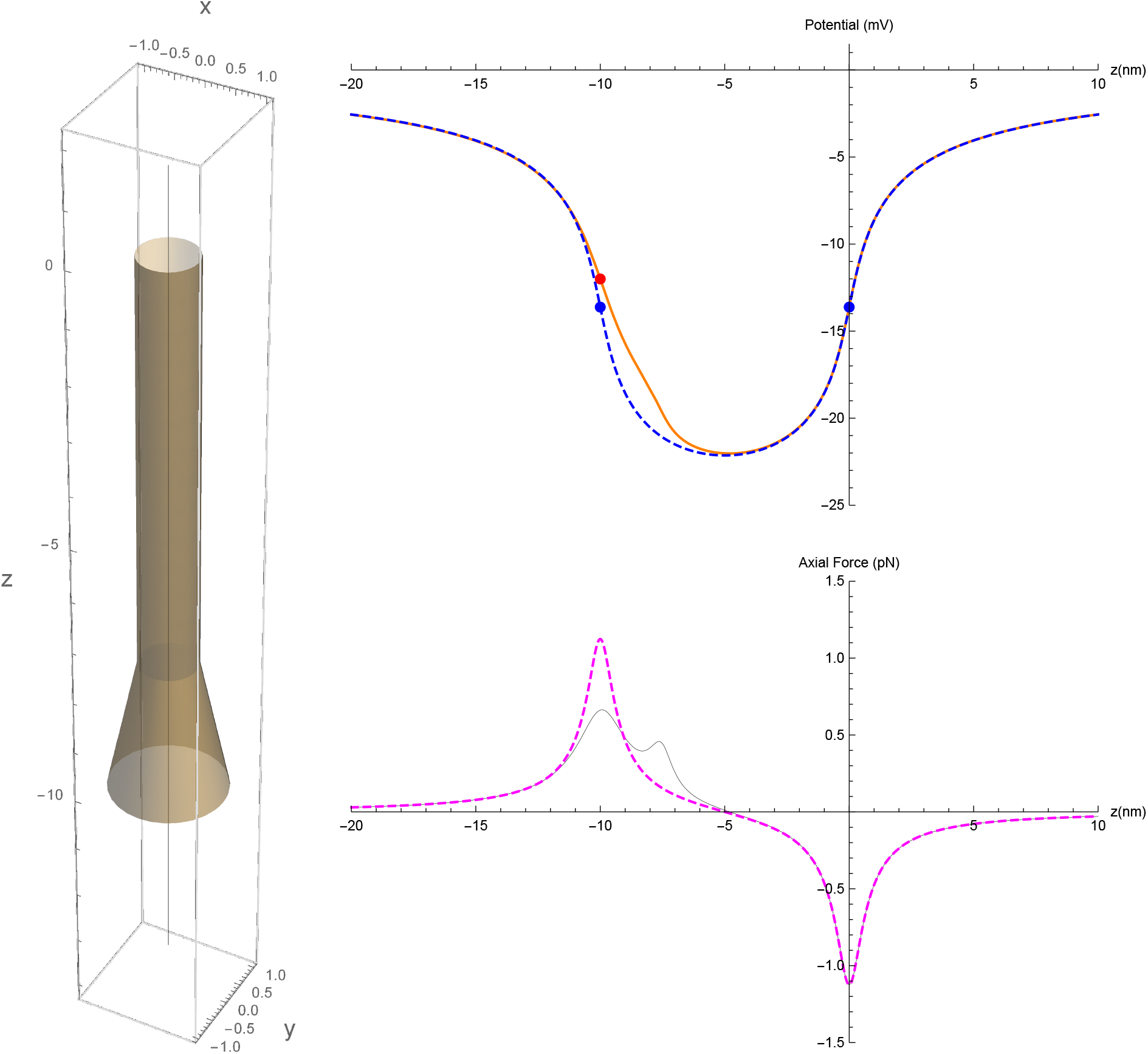
Electrostatic scalar potential (upper panel) and axial force (lower panel) profiles for a hollow cylinder concatenated to a truncated cone. The transition between the cylinder and the truncated cone is at *λ* = 0.75 (see text). Dashed lines: cylinder only; full lines: cylinder concatenated to cone. Red and blue points: the potential at the tunnel exit point is higher than the potential at the entry point. The axial forces in the tunnel exit region are smoother and more dispersed in the combined geometry.

This is unfavorable to the positively charged amino acid residues that will have to traverse the tunnel and will require more mechanical energy to overcome this electrostatic potential difference than their negatively charged amino acid counterparts, when moving from the entry point *z* = 0 to the exit point *z* = −*L*. In an adopted geometry that best fits the experimental observations (see below), with *λ* = 0.75*, α* = 0.198 and with *σ*_2_ = 2/3 *σ*_1_, the potential difference is −12.65 − (−14.35) = 1.70 mV and the required mechanical work for traversing the tunnel is 0.164 kJ/mol ~ 0.039 kcal/mol for a positive unit charge embedded in an otherwise neutral nascent peptide stretch.

This provides a rough estimate of the energy required for a single positively charged amino acid residue to traverse the ribosomal exit tunnel if this positive residue is embedded in a completely neutral peptide sequence. The mechanical energy requirement for real sequences depends on the particular distribution of the charged amino acid residues along the primary sequence. There might be particular sequence contexts for which the local mechanical energy requirements could be much higher than the estimated values given above.

For a straight cylinder, the mechanical energy is equal to zero (symmetry in the potential between exit and entry points), whereas for the truncated cone concatenated to the cylinder (asymmetry in the potential between the exit and entry points), the mechanical energy uptake when moving a single positive unit charge from the entry point (*z* = 0) to the exit point (*z* = −*L*) is estimated to be around 164 Nm/mol for a stretch of 40 residues in the tunnel, according to our electrostatic interaction model. The estimated mechanical energy uptake per residue incorporation would be around 4.1 Nm/mol per residue incorporation(~ 0.001 kcal/mol per residue incorporation).

This is due to the fact that the axial forces profile is not symmetrical in the cone concatenated to the cylinder geometry. However, in this truncated cone geometry (asymmetrical potential), the axial forces amplitudes are reduced and are more spatially dispersed than in the cylindrical geometry as displayed in Fig. 5.

### D. Permittivity of the medium prevailing in the tunnel

The medium inside the ribosome wall is of course a dielectric and not a conductor and not vacuum. In formula (19), we see that the potential on the cylinder axis depends on the geometry and surface charge density. The *ϵ*_0_ parameter is the vacuum’s permittivity: 8.854 10^−12^ Farad/m. The formula should be generalized to apply to the real dielectric aqueous medium prevailing inside the ribosome exit tunnel. The relative permittivity of water is *ϵ*_*r*_ = 78 at 25 Celsius degrees and around *ϵ*_*r*_ = 74 at 37 Celsius degrees. So, the vacuum permittivity should be replaced by a more appropriate dielectric permittivity: *ϵ* = *ϵ*_0_ · *ϵ*_*r*_ where *ϵ*_*r*_ = 74. The presence of specific water soluble free cations such as Mg^2+^ or Zn^2+^ may also change the medium permittivity inside the tunnel. The ionic strength inside the tunnel and the electric polarizability of all the molecules inside the tunnel would of course also play a role that is neglected here in our simplified model.

Simple volume calculations show that the number of water molecules occupying the inner volume of the ribosome exit tunnel is of the order ~ 1, 000 when no polypeptide is present. This is considered large enough for a continuous classical physical theory to still be applicable. At smaller scales, quantum effects or atomistic molecular dynamics effects should be incorporated. Lucent *et al.* [40] indeed advocated that the understanding of the complexity of molecular behavior in the ribosome exit tunnel should require an atomistic description including the solvent confined to the tunnel. Their simulations showed that solvent (water) confined to the tunnel does not behave as a continuous isotropic dielectric medium. In our simplified approach, we nevertheless considered the isotropic dielectric permittivity of the tunnel medium to be the one of water as a bulk-like solvent.

### E. Estimation of the number of negatively charged phosphate groups exposed on the tunnel inner wall and of the surface charge density of the ribosome tunnel inner wall

In eukaryotes, one of the rRNA molecules in the 60S ribosome subunit is the 28S rRNA. The 28S rRNA size is 5,034 nucleotides in higher eukaryotes (*H. sapiens*) and entails 5,034 phosphate groups. It is not known how many phosphate groups are really exposed inward the tunnel. There is not a net negative charge on each phosphate group but rather, each phosphate group has a formal negative charge that would instead be a small fraction of a net negative charge. As a first wild guess, we assumed there are 1,000 exposed groups and that the formal negative charge distributed on the oxygen atoms of the phosphate group is such that each phosphate group actually carries a partial charge representing a fraction of 1 over 1,000 unit charge. Then the net negative charges lining the surface of the inner wall of the tunnel would sum up to −1 times the unit charge. This is equivalent to a single net negative unit charge for the whole tunnel inner surface.

Given the electrostatic potential values that have been experimentally measured by Carol Deutsch and co-workers [29], we can get an estimate of the surface charge density of the ribosome inner wall. Indeed, for a measured potential in the range −15 mV to −22 mV, and assuming the geometry of the cylinder is *L* = 10 nm and *R* = 0.5 nm, substituting *ϵ*_0_ with *ϵ* = *ϵ*_0_ · *ϵ*_*r*_, an estimate for *σ* using formula (19) is −10.2 10^−3^ *C/m*^2^, if the relative permittivity of water at 37 Celsius degrees is adopted for the ribosome exit tunnel.

The cylinder surface being 3.14 10^−17^m^2^, the total apparent surface charge would be around −3.2 10^−19^C on the tunnel inner wall. Dividing by the unit charge of an electron, i.e. 1.602 10^−19^ C, the number of net unit charges would be around 2 and the charged fraction of the phosphate groups would be around 0.002 (under the previous assumption of 1, 000 exposed phosphate groups), indeed complying with the fact that the atoms are actually partially charged and reflecting the electronic cloud density distribution in the molecular structure of the phosphate moiety esterified to the ribose sugars in the rRNA backbone.

## III. REALISTIC ELECTROSTATIC MODELS OF THE RIBOSOME EXIT TUNNEL

### A. Realistic electrostatic potential profile of cylindrical shape best fitted to experimental point measurements

The real electrostatic potential profile inside the ribosome exit tunnel was experimentally measured with an ingenious biochemical technique of molecular tape at least in prokaryotic ribosomes by Carol Deutsch and co-workers [29]. To our knowledge, the potential profile in eukaryotic ribosomes exit tunnels has not been measured experimentally yet. The real electrostatic potential profile is actually not symmetric. We further need to build an improved and more realistic potential profile by adding to the previous idealized models a small Lorentzian peak function. The motivation for this comes from the experimental data showing that the electrostatic potential locally increases at a distance one third of the length of the tunnel away from the PTC center (approximately at least 15-17 amino acid residues in the nascent protein upstream from the amino acid residue incorporation site). This local increase in the potential is located near the position of the ribosomal constriction, where specific ribosomal conserved constitutive proteins protrude inward the tunnel, i.e. L4 both in bacteria and eukaryotes, L22 in bacteria and L17 in eukaryotes, see Fig. 1 (left). Dao Duc *et al.* [38] confirmed, with multiple sequence alignments of uL22 and L4 proteins across 20 species in the three domain of life, the presence of a highly conserved sequence enriched in arginine (R) and or lysine (K). In uL22, there are up to 7 R or Ks conserved between position 154 and 176. In L4, there are 5 Rs (or Ks) conserved between position 71 and 92 across eukaryotic species and up to 6 Ks or Rs conserved between position 69 and 82 across prokaryotic species. Similar conservation has been shown for uL23 (bacteria) and eL39 (eukaryotes) [38]. These positively charged residues protrude near the tunnel constriction and explain the local rise of the potential. The Lorentzian local peak potential as expressed in equation (45) that we added was fitted to the experimental data obtained by Lu *et al.* [29]

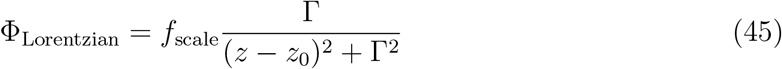

The fitted parameters values are *f*_scale_ = 9 10^−9^ Volt · m for the scale factor, Γ = 9 10^−10^ m for the Lorentzian peak full width at half maximum and *z*_0_ = −3.75 10^−9^ m for the peak center location, i.e. 37.5Å measured from the entry side point towards the protein tunnel exit. The experimental data points taken from Lu *et al*. [29] and the fitted adapted function for the ribosome exit tunnel electrostatic potential are displayed on the upper panel of Fig. 6 when the simple straight cylinder geometry is adopted. The extended expression for the total electrostatic potential in this improved model version is

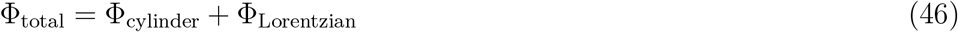

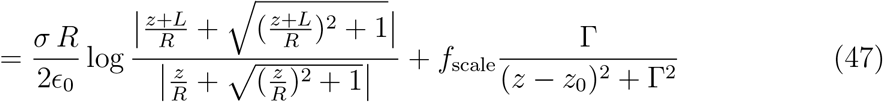

**FIG. 6:**
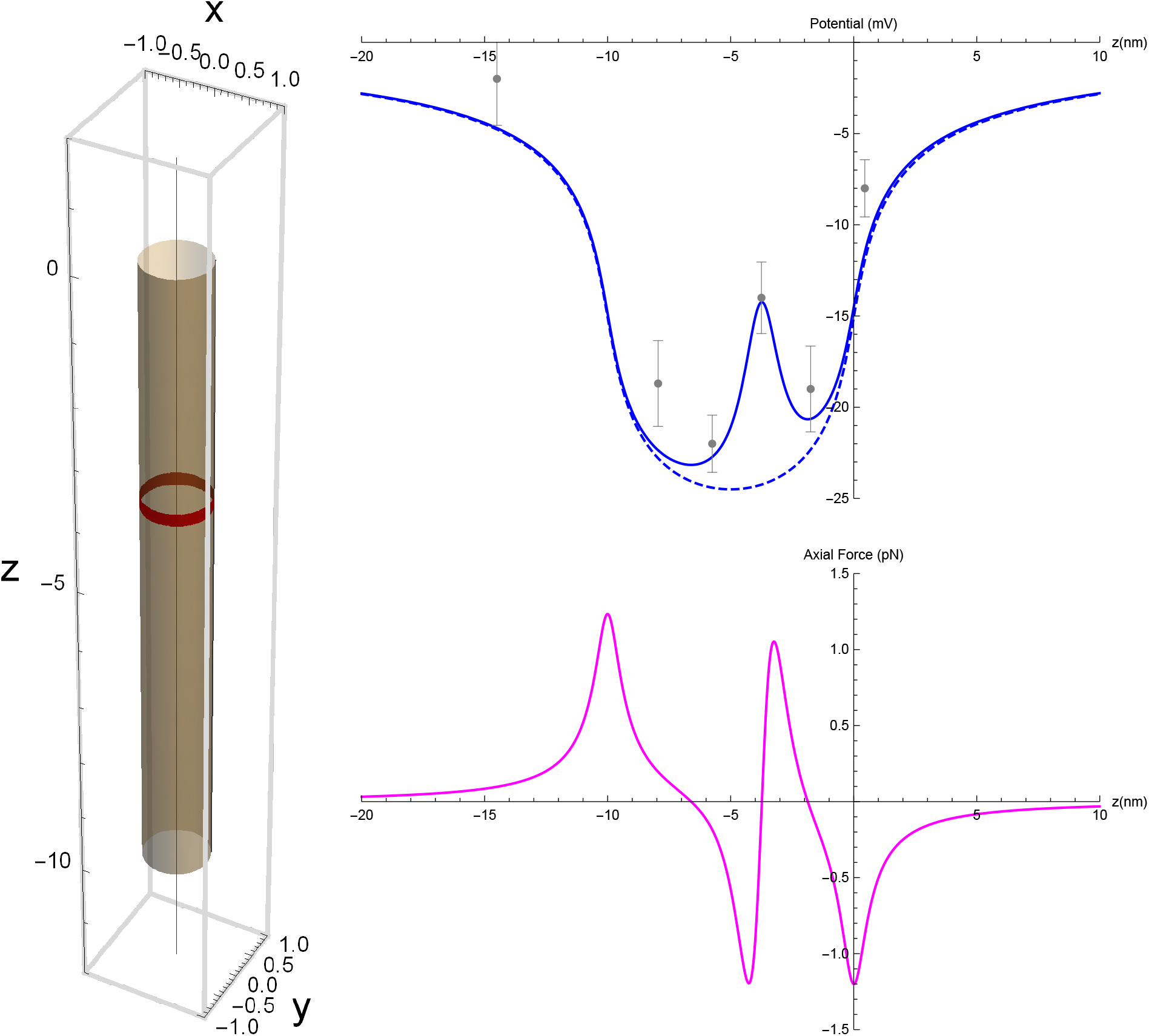
Improved model for the electrostatic scalar potential of the ribosomal exit tunnel. A Lorentzian peak was added locally to the idealized cylindrical geometry potential which was fitted to the experimental data points obtained by Lu *et al.* [29] This local peak accounts for the ribosomal protein protrusion (L22 or L17, L4) inside the tunnel (upper panel). The protrusion’s position is indicated by the red ring (left panel). Dashed line: potential resulting from the idealized uniformly charged hollow cylinder. 95% confidence intervals error bars computed from the experimental data. The axial forces for a positively unit charged test amino acid residue on the tunnel axis as a function of axial position in the tunnel for the improved model (lower panel).

An important characteristic of the Lorentzian function Φ_Lorentzian_ that is shared with the potential Φ_cylinder_ is its vanishing at infinity in both directions.

The total electric field (and the force) is obtained by

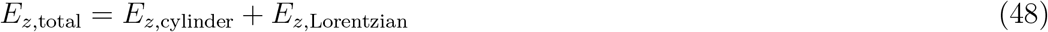

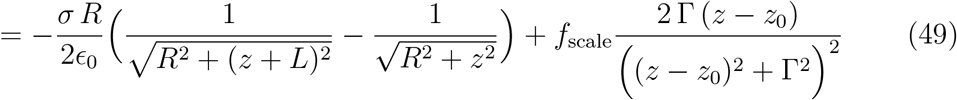

from which the axial force results immediately by *F_z_* = *q* · *E_z_* and is displayed in Fig. 6 lower panel in the case of a single positively unit charged amino acid residue.

The charge surface density *σ* in formula (19) could also be made dependent on the *z* variable to account for local heterogeneity on the ribosome wall and to account for the experimentally observed potential profile.

### B. Realistic electrostatic potential profile of truncated cone combined to cylindrical shape best fitted to experimental point measurements

A still better fit of the experimental data of Deutsch and co-workers [29] is obtained with the truncated cone concatenated to the cylinder geometry. Keeping the same Lorentzian peak, the best extended expression for the total electrostatic potential in this last improved version of the model is

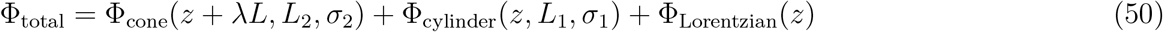

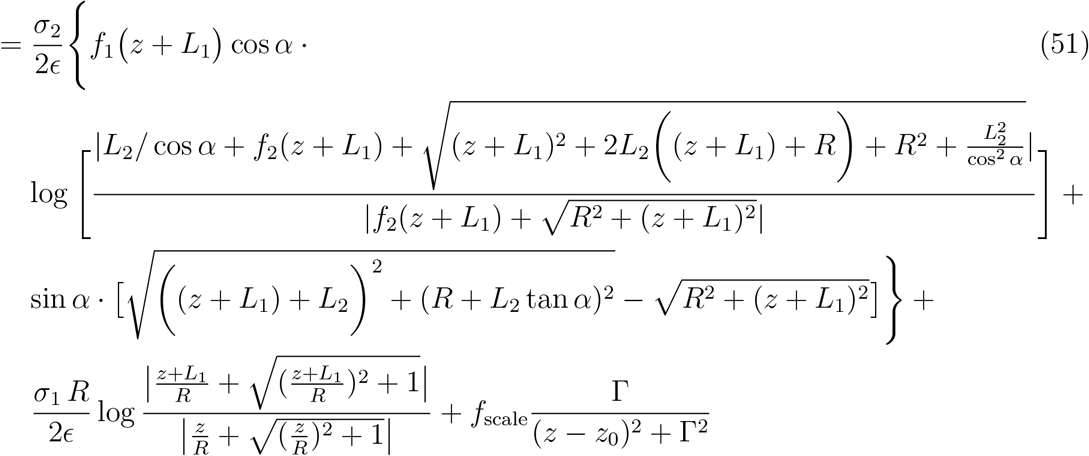

where *L*_1_ = *λ*·*L*, *L*_2_ = (1−*λ*)·*L*, *L* = *L*_1_ +*L*_2_ and *σ*_1_*, σ*_2_ comply with the charge conservation that was exposed previously and consistent with the assumption of an elastic deformation of a preexisting cylinder surface turned into a truncated cone surface preserving the same total charge.

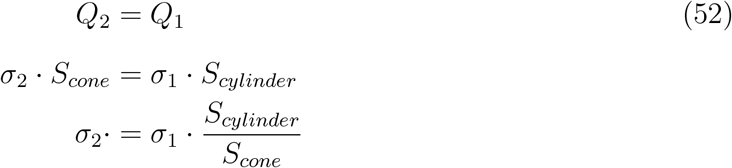

Similarly for the axial electric field (and the axial force) along the *z*−axis

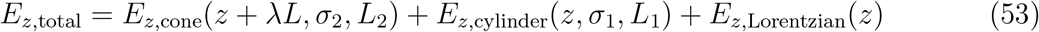

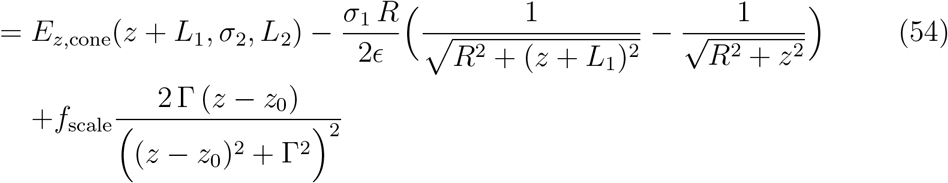

where the detailed expression of the first term in the last right hand side is easily obtained from equation (41) by substituting *z* + *λL* to L, *σ*_2_ to *σ* and *L*_2_ to *L*. The plots of the electrostatic potential and of the total axial force profiles in the ribosome exit tunnel under this last improvement of the model are displayed in Fig. 7. The upper panel shows the goodness of the fit with the experimental data of Lu *et al.* [29] The improvement of the fit due to the cone geometry concatenated to the hollow cylinder is worth noticing (orange line in the upper panel of Fig. 7). This last version of the model perfectly fits the four experimental points located inside the tunnel.

**FIG. 7:**
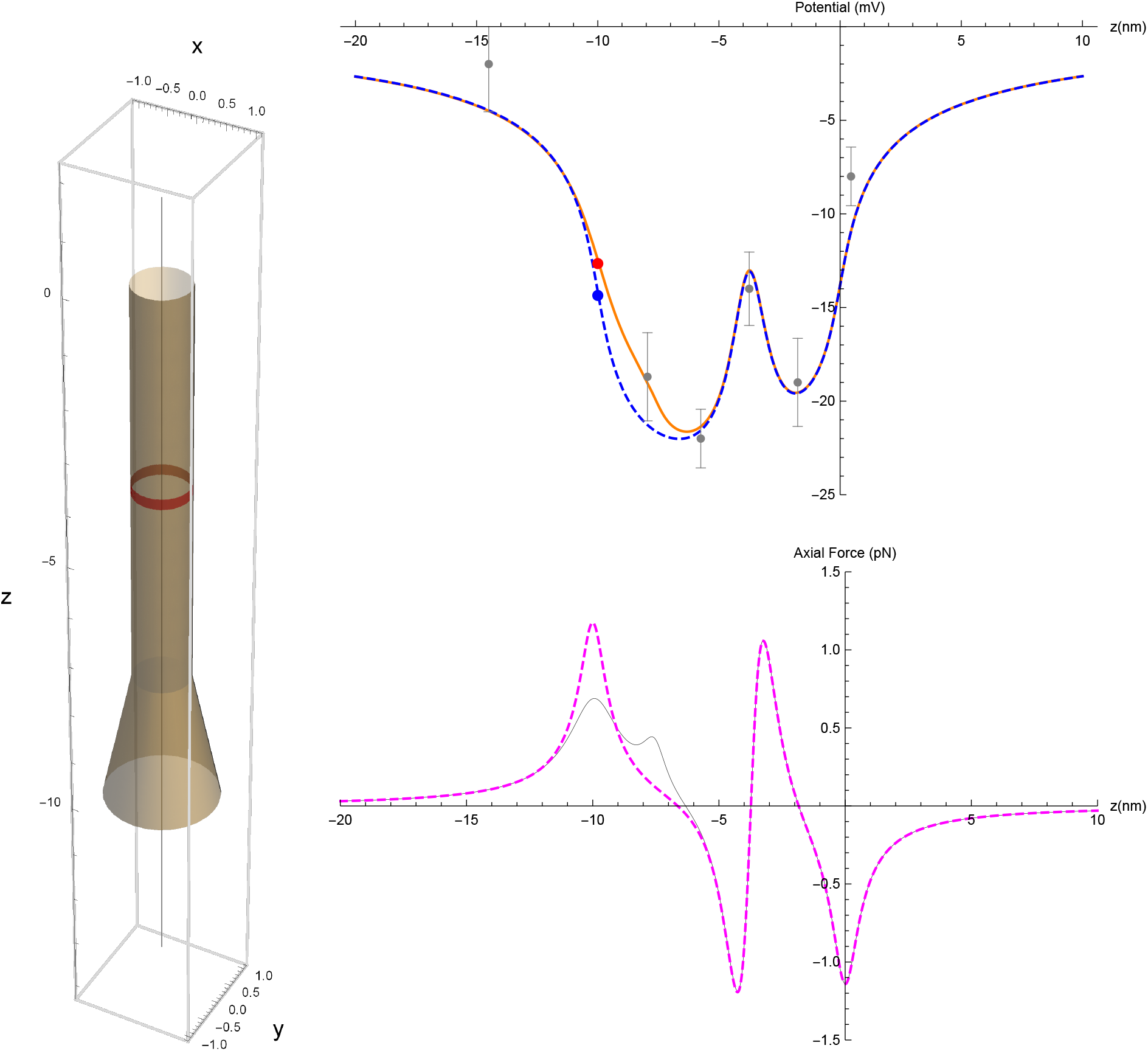
Best fitted model for the electrostatic scalar potential of the ribosomal exit tunnel. Upper panel: a Lorentzian peak was added locally to the idealized conical plus cylindrical geometry potential which was fitted to the experimental data points obtained by Lu *et al.* [29]. The protein protrusion’s position is indicated by the red ring (left panel). Dashed blue line: potential resulting from the idealized uniformly charged hollow cylinder. Orange line: potential resulting from the superposition of a cylinder and a truncated cone preserving the same total charge. 95% confidence intervals error bars computed from the experimental data. Lower panel: the axial forces for a positively unit charged test amino acid residue on the tunnel axis as a function of axial position in the tunnel for the best fitted model. Black line: axial forces profile resulting from the superposition of a cylinder and a truncated cone. Dashed magenta: idealized uniformly charged hollow cylinder.

The model is a very good fit of all the 6 experimental data points if the tunnel length is taken in a range from *L* = 8.5 nm to *L* = 9.5 nm, keeping all the other parameters constant. Fig. 8 shows the plots for the potential curves for these two tunnel lengths boundaries and shows that the 6 experimental measurements are correctly captured within the 95% confidence intervals of the potential measurements between these two length boundaries. In their study, Lu *et al.* mapped their 6 experimental points on the ribosomal crystal structure of *Haloarcula marismortui* (archae) for which the cryo-electron microscopy resolved ribosome structure gives a tunnel length of ~ 9.5 nm [38]. It is recognized that there might be some deviations in mapping distance and with respect to the actual length of the ribosome exit tunnel of the biological material that they used. Also, the actual in vivo lengths might slightly differ from the lengths determined in the cryogenic conditions prevailing in cryoelectron microscopy and the functional length might also slightly differ to the geometrical length. The cryo-electron microscopy ribosome structure resolution conducted on 23 species across the three domain of life are supporting the fact that the ribosome exit tunnel in bacteria is a bit longer than the one in eukaryotes while archea have intermediate lengths between bacteria and eukaryotes [38]. Hence, a tunnel length of *L* ≈ 8.5 nm, should be adopted if we aim at building a model for the eukaryotic/mammalian ribosome exit tunnel, that could be eventually used for computational biology and bioinformatics purposes on eukaryotic/mammalian omics data. The plot of the electrostatic potential in a ribosome exit tunnel with a length of 8.5 nm and with a variable angle of the cone frustum at the tunnel exit is shown in the animated figure Fig. S1, provided in the supplemental material.

**FIG. 8:**
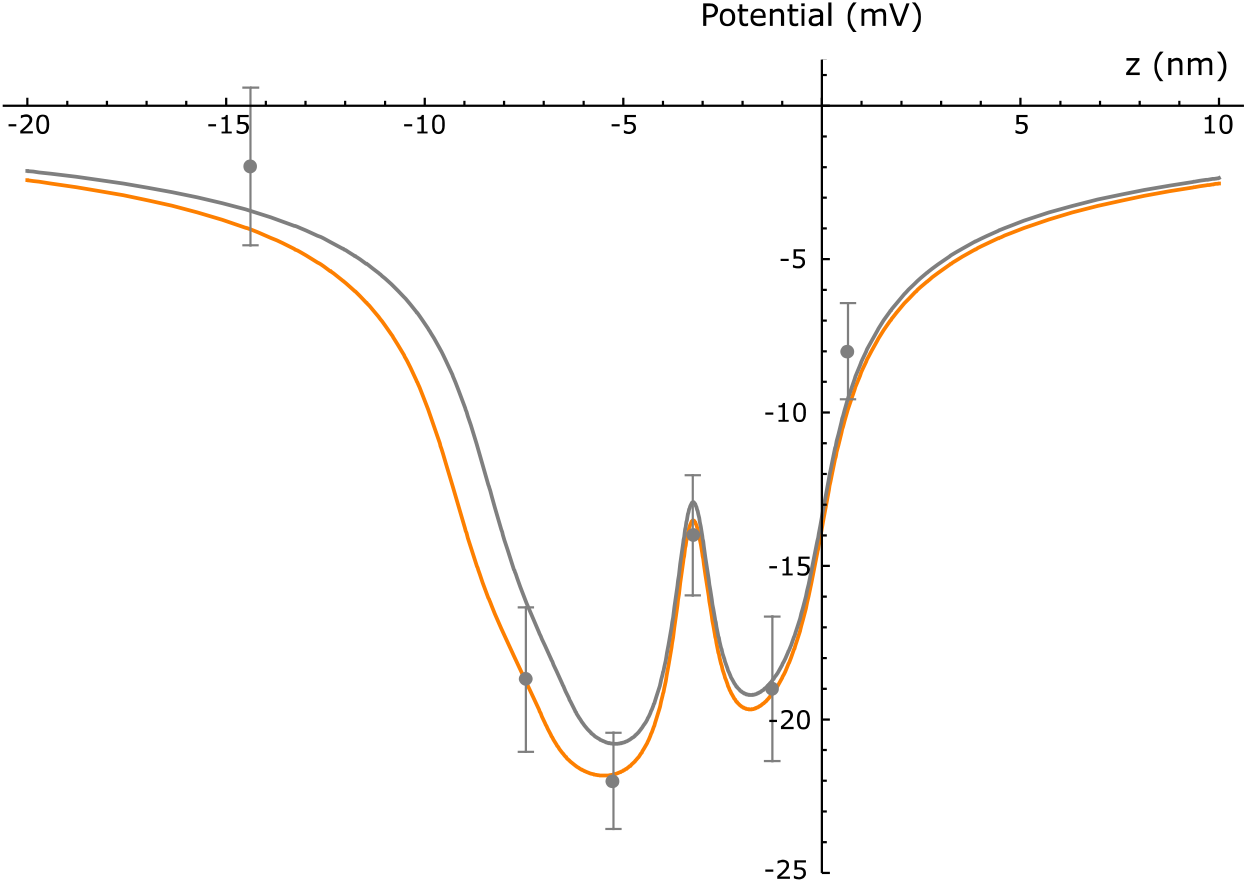
Electrostatic scalar potential curves for two tunnel length boundaries. Tunnel length = 9.5 nm: orange curve. Tunnel length = 8.5 nm: gray curve. Both curves capture the 6 experimental points within the 95% confidence interval of the potential measurements that were mapped on the ribosomal crystal structure of *H. marismortui* by Lu *et al* [29].

### C. Electrostatic potential modulation due to dynamical change in the geometry of the exit tunnel

The ribosome is a complex molecular machine [41]. The dynamical conformational changes in the ribosome during codon translocation have also been extensively investigated for the last two decades. The two subunits of the ribosome (small subunit SSU and large subunit LSU) are moving relative to each other [42] and there are also evidence of conformational changes internal to the small subunit during tRNA recognition and accommodation (swiveling motion of the SSU subdomain, the SSU head, relative to the SSU body) [1, 42, 43]. Under these new insights into the ribosome dynamics during translation, it is important to take into account the internal forces that could arise from these geometrical changes, particularly when fixed formal electrical charges are spatially moved relatively to one another. Our electrostatic model of the nascent chain exit tunnel in the large subunit LSU of the ribosome shows there are important physical effects associated to such conformational changes in the shape of the tunnel with potent functional and biological consequences.

Changes in the intensity of fluorescence resonance energy transfer (FRET) between fluorescence reporters attached in pairs at different SSU and LSU positions on ribosomes have shown that during translation and for each translocation step occurring at codon reading and amino acid incorporation in the nascent peptide, a relative rotation of the ribosome large subunit with respect to the small subunit occurs [41–43].

Although, to our knowledge, this has never been investigated experimentally, we explore here, from a theoretical perspective, the possibility of a reversible elastic deformation in the exit tunnel geometry in the large ribosomal subunit (LSU). Indeed, it should not be taken for granted that the tunnel geometry is constant over time. An enlargement or a narrowing of the cone frustrum at the tunnel exit with conservation of the electric charge supported by the inner wall of the tunnel would cause a modulation of the electrostatic potential inside the tunnel and would exert a change in the net axial force on the charged amino acid residues that are in the nascent protein stretch locally occupying the tunnel. To give a pictorial view of this dynamic conformational change, an animated figure is provided in the supplemental materials showing how a change in the tunnel geometry changes the electrostatic potential inside the tunnel dynamically (see Fig. S1). According to this proposed hypothesis, the ribosome exit tunnel could be viewed as an electrostatic braking or pushing device helping the nascent protein to either progress through the tunnel or to fold properly, thereby contributing to the translational control of protein synthesis. The molecular basis of these dynamical conformational changes in the exit tunnel shape is unknown.

## IV. APPLICATION OF THE RIBOSOME EXIT TUNNEL MODEL

### A. Computing the electrostatic interaction variations of the ribosome exit tunnel for different amino acid sequences in nascent polypeptides

Our established models expressed by equation (19) or (47) or (51) can be used to quantitatively compare how difficult it is for the ribosome to push a nascent polypeptide chain inside and eventually out of the exit tunnel, depending on the amino acid primary sequence. If a peptide sequence is locally enriched in positively charged amino acids residues inside the tunnel and in negatively charged amino acid residues close to tunnel entry point, the axial forces required to push the nascent protein through the tunnel will be higher than for a peptide composed of neutral amino acid residues in the primary sequence or carrying only a single cluster of positively charged amino acid residues. The electrostatic potential well, locally trapping charged amino acid enriched peptides, needs to be overcome by other forces. These compensation mechanisms are exerted either by the ribosome itself or by third party proteins with motor domains from specialized chaperone proteins exerting tugging forces outside of the ribosome. The elongation speed also has to be compatible with the decoding speed of the mRNA encrypted message which depends on the codon usage and on codon position autocorrelation, i.e. codon ordering allowing tRNA recycling (reusage of the same tRNA at successive encodings of the same amino acid can speed up translation or favor fidelity [44, 45]). The elongation speed may also independently be impeded by downstream mRNA secondary structures [21, 22, 33, 34].

The eukaryote ribosome exit tunnel can accommodate at least 40 amino acid residues and up to more than 70. It is known that the nascent polypeptide can start folding, i.e. finding its final secondary structure, inside the tunnel (and eventually tertiary 3D structure outside). Alpha helices secondary structures have been shown to be present inside the tunnel close to its exit point. So, a variable number of amino acids larger than 40 can actually be hosted inside the tunnel. Again, for the sake of simplicity, here, we consider that the number of amino acid residues hosted inside the tunnel is exactly 40 and that the maximum number of amino acid residues that are under the electrostatic influence of the tunnel is exactly 50: 5 between the PTC and the tunnel entry, 40 in the tunnel and 5 out of the tunnel.

The incorporation of a single amino acid to the nascent polypeptide chain takes place at the peptidyl transfer center (PTC), at the so called P site of a translating ribosome. This PTC center is located around 5 amino acid residues away from the ribosome tunnel entry point. Stated otherwise, this means that the currently decoded codon, for which the cognate or semi-cognate aminoacylated tRNA, is 5 codons downstream the codon for which the amino acid is currently in the entry point of the tunnel. There are 5 amino acids bound in the oligopeptide part ready to enter the tunnel. Let us also assume that there are 5 bound amino acids out of the tunnel at the exit side that can feel the electrostatic influence of the tunnel. So, from the start codon (AUG coding for methionine), a nascent peptide starts with a 5 amino acid residues stretch elongating to the tunnel entry point, building up progressively to a 45 amino acids sequence fully accommodating the whole length of the tunnel, eventually extending to 50 amino acid residues being under a direct influence of the tunnel, see Fig. 9. For this 50−mer stretch to be out of the tunnel influence, another extra 50 amino acids have to be added to the carboxy terminal end of the nascent polypeptide.

**FIG. 9:**
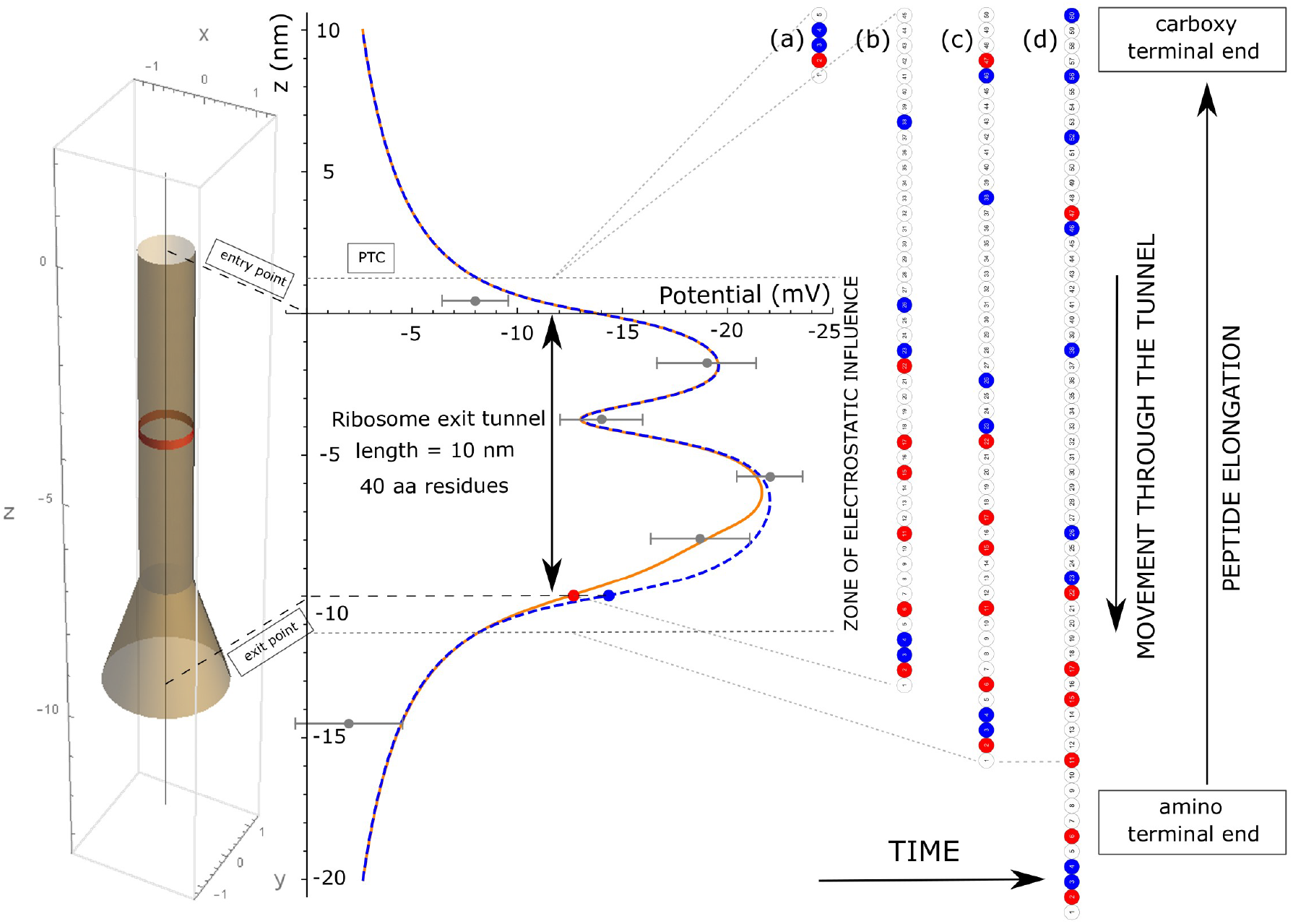
Algorithm for computing the axial forces acting on a nascent peptide. Only the last 50 residues, at most, from the PTC are under the electrostatic influence of the ribosome exit tunnel. In this figure, the nascent polypeptide is moving from the top to the bottom through the ribosome exit tunnel during elongation.(a) 5 amino acid residues peptide with residue 5 at the PTC and residue 1 at the tunnel entry point. (b) 45 amino acid residues with residue 45 at the PTC and residue 1 at the tunnel exit point. (c) 50 amino acid residues with residue 50 at the PTC and residue 1 emerging out of the electrostatic influence zone. (d) 60 amino acid residues with residue 60 at the PTC and residues 1 to 10 out of the electrostatic influence zone.

Our aim is to compute the force profile and the mechanical power to be applied continuously on peptide stretches to overcome the electrostatic trapping interaction in the tunnel and to exit the ribosome tunnel. The easiest case scenario for computing would be when the downstream sequence is completely neutral. This is of course not always the case and the occurrence of charges in the downstream sequence plays a role that should not be neglected.

Multiplying the axial force acting on the stretches with the stretch axial displacement, i.e. the elongation distance towards the ribosome exit tunnel, yields the mechanical work that was delivered. Multiplying the axial forces acting on the stretches with the protein elongation rate, i.e. the speed of the ribosome along the transcript (mRNA) being translated, yields the required instantaneous net mechanical power.

The electrostatic axial force on the tunnel axis felt by an amino acid residue is the product of its net charge by the axial electric field, see equation (23), the latter being the gradient of the electrostatic scalar potential, i.e. the first derivative of the potential with respect to the axial coordinate.

An important simplifying assumption is that all amino acid residues building up the nascent polypeptide are all rigidly bound together and that the resulting nascent protein can be considered a single linear solid rigid body. This peptide, at least in the tunnel, is considered non deformable. With this strong assumption, the axial forces individually computed for each charged amino acids act jointly and apply additively on the resulting rigid peptide body.

The local pH along the tunnel is unlikely to be out of the range 6-8 [46]. In this pH range, among the 20 amino acid residues, only three are positively charged and two are negatively charged. Arginine (R), lysine (K) and histidine (H) carry a partial positive charge on the amino moiety in the side chain. The intrinsic pK value, referred to as pK_int_, is the pK value of an ionizable side chain when it is present in pentapeptides [30]. Only arginine, pK_int_ = 12.3, and lysine, pK_int_ = 10.4, are truly positive in physiological conditions at neutral pH whereas histidine, pK_int_ = 6.5, would be very weakly positive at a pH in the range 6 − 6.5. For this reason +1, +1, +0.05 net formal charges are arbitrarily adopted for arginine, lysine and histidine respectively. For glutamate (E), pK_int_ = 4.3 and aspartate (D), pK_int_ = 3.9, both carrying a carboxylic moiety on the side chain, the arbitrarily adopted net formal charges are both −1 in physiological conditions. All other amino acid residues are considered neutral. The positively charged residues are represented in red whereas negatively charged residues are represented in blue on the test sequences to be analyzed under our model as displayed in Fig 9. The neutral residues are unsensitive to the electrostatic potential or the axial electric field.

#### Algorithm and program pseudo-code for computing the axial force on the nascent peptide due to the ribosome exit tunnel interaction

The algorithm for computing the axial force on a given nascent peptide due to the ribosome exit tunnel electrostatic interaction as a function of the amino acid sequence is schematically depicted in Fig. 9.

#### Reading in the given input peptide sequence

(Step a) Read in the peptide sequence from the amino terminal end to the carboxy terminal end.

(Step b) Determine the length of the peptide (number of amino acid residues in the given peptide).

(Step c) Convert the sequence of amino acid residues into an ordered list of formal charges using the following charge coding rule: *K* → +1*, R* → +1*, H* → +0.05*, E* → −1*, D* → −1. All other residues are converted to a neutral charge *X* → 0.

#### Computing the axial position of each amino acid in the sequence, compute the axial force acting on the residue at that position and sum the contributions of all charged residues

(Step a) Start with the first 5 residues from the amino terminal end of the peptide (the first five elements in the ordered list) to build the stretch currently computed.

(Step b) Map the axial positions of the residues in the stretch, each separated by a distance 0.25 10^−9^ m. Position *z* = 0 corresponds to the residue located at the ribosome exit tunnel entry point, position *z* = 5 × 0.25 10^−9^ corresponds to the residue located at the PTC, 5 residues downstream in the sequence. All algebraic negative *z* positions correspond to residues that have entered the tunnel.

(Step c) Compute the ordered list of axial electric fields for each of the previous axial positions using formula (22) for the idealized cylindrical model, formula (49) or formula (55) for the realistic model, incorporating the Lorentzian peak and the truncated cone geometry at the end side of the tunnel, respectively.

(Step d) Multiply element by element, the ordered list of the axial electric fields by the ordered list of formal charges, to obtain the list of the contributing axial forces acting on the peptide stretch currently computed.

(Step e) Sum all the contributing axial forces in the peptide stretch currently computed and store the result in an ordered list of the total axial forces acting on the stretch from the PTC site.

(Step f) Repeat *Step b* to *Step e* for all iterated stretches by one residue towards the carboxy terminal end, conditionally on a length of 50 residues, and while the last residue has not reached the end of the given input peptide. The 50 residues condition ensures there are at most 40 residues inside the tunnel, 5 residues between the PTC site and the tunnel entry point and at most 5 outside the ribosome exit tunnel, still under the electrostatic influence of the tunnel.

#### Plot the total axial force acting on the nascent peptide as a function of the last amino acid residue occupying the ribosomal PTC position

Positive axial forces are believed to slow down the elongation rate while negative axial forces are believed to speed up the elongation rate of the ribosome.

### B. Comparing the electrostatic interaction profiles when passing through the ribosome exit tunnel for different amino acid sequences

#### 1. Simulated synthetic oligopeptide sequences

It should be emphasized that due to the symmetry of the potential barrier in the idealized cylindrical model and its finite length, a clustered local enrichment in positive (negative) charge in a polypeptide sequence will first be attracted (repelled) when entering into the tunnel and will then be pulled inside (pushed outside) the tunnel when emerging at the tunnel exit point. Hence an inversion in the sign of the force profile should always be observed for locally clustered net charges that are followed by a neutral tail sequence. This inversion spreads over a distance covering the ribosome exit tunnel length which is 40 amino acid residue in length in the adopted simplified model and with equal areas under the curve, see Fig. 10 (A) and (C).

**FIG. 10:**
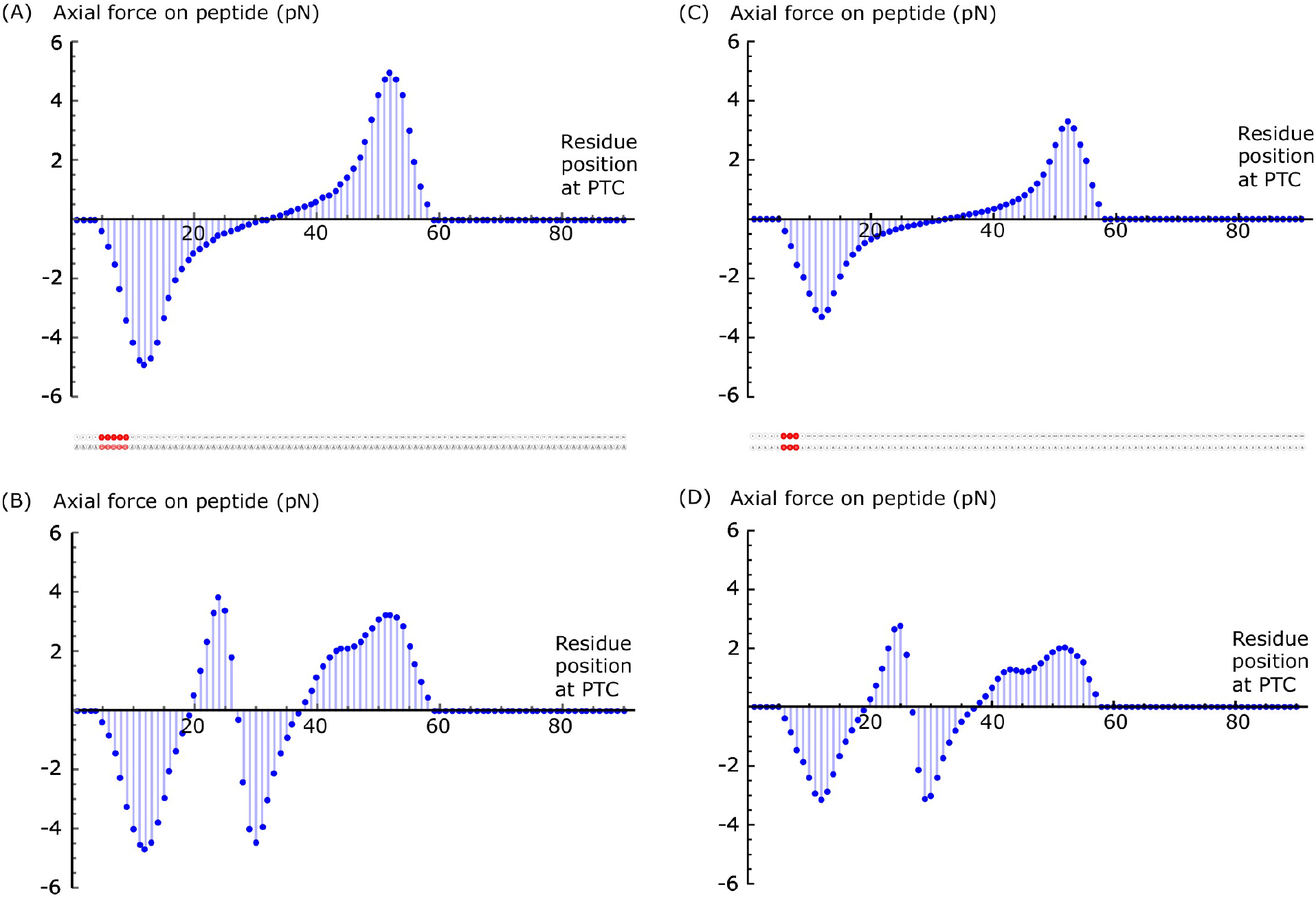
Axial forces (pN) ordered values acting on the nascent chain at each residue position incorporated in the primary sequence at the PTC center. Symmetric electrostatic potential idealized model (A) with 5, (C) with 3 contiguous arginine. Asymmetric electrostatic potential realistic model (B) with 5, (D) with 3 contiguous arginine, centered at position 7. Positively charged arginine residues are colored in red in the primary sequence as displayed in the figure insets.

The situation is more complicated when the tail sequence also includes local charges distribution within a range of 20 − 40 amino acid residues in the tail sequence or if the electrostatic potential well barrier is not symmetric as with the truncated cone concatenated to the cylinder geometry.

To highlight the differences between a symmetric potential (idealized cylindrical model) and an asymmetric potential (cylinder plus truncated cone with Lorentzian peak realistic model), we compared the axial force profiles applied for the same synthetic sequences in both cases with typical clustered net charge distributions.

In Fig. 10 to 11, the axial forces acting after each amino acid incorporation at the PTC are displayed for a peptide of 90 residues in length. Fig. 10 (A) shows the symmetric potential (idealized cylindrical model) effect on 5 contiguous positively charged residues between position 5 and 9 (net positive charge centered at position 7). The nascent peptide is attracted into the tunnel until amino acid residue number 32 (= 7+25) is incorporated at the PTC. From position 32 to 59 = 9 + 50 (position 59 corresponds to the moment when the last positively charged residue is out of the influence zone), the axial forces acting on the peptide tend to pull it back into the tunnel and these forces tend to prevent the peptide from traversing the tunnel easily.

**FIG. 11:**
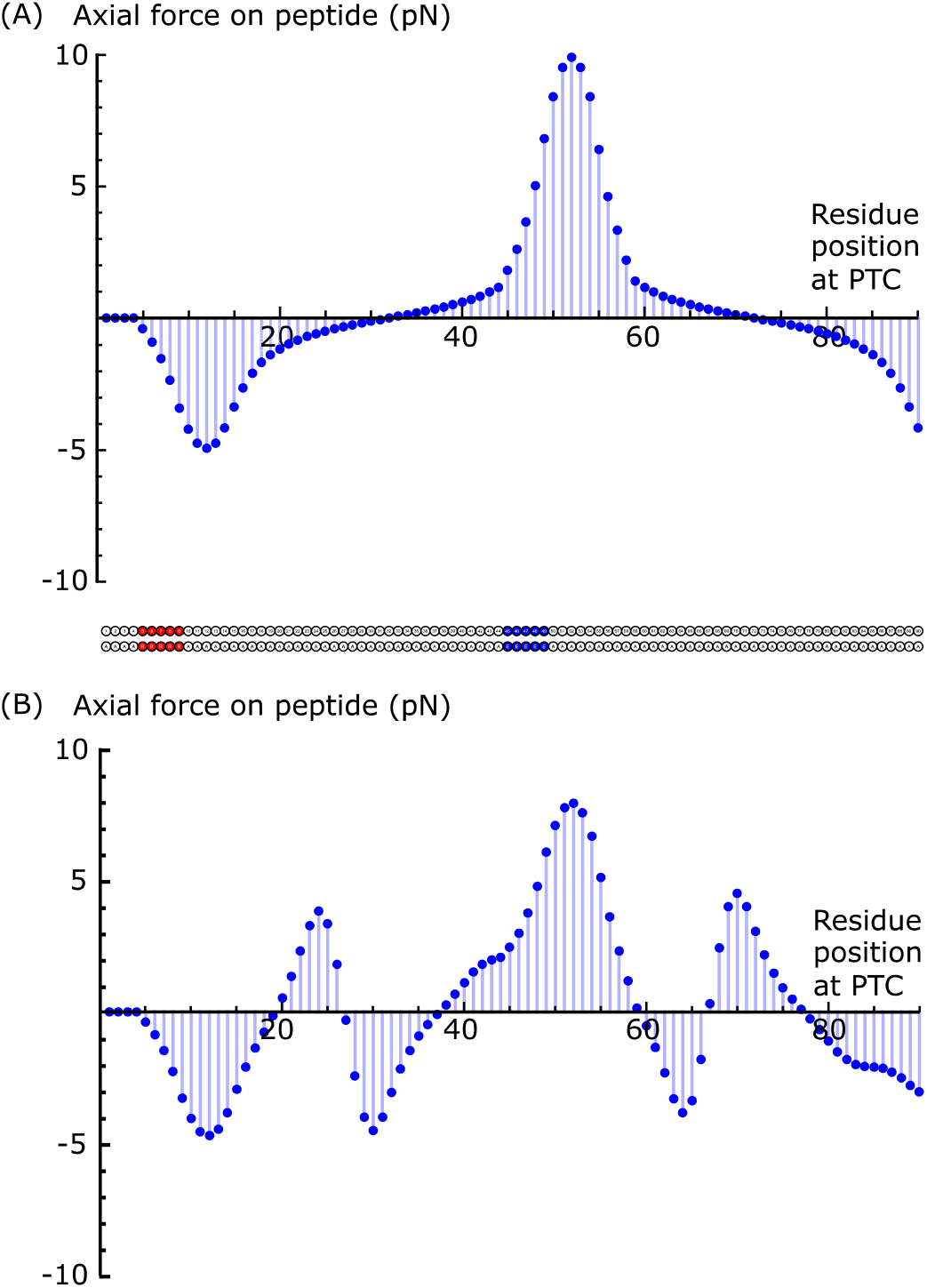
Axial forces (pN) ordered values acting on the nascent chain at each residue position incorporated in the primary sequence at the PTC center. Symmetric electrostatic potential idealized model (A) with 5 contiguous arginine and 5 glutamate residues clustered as displayed in the figure inset. (B) Asymmetric electrostatic potential realistic model. Opposite charges (+ residues position 5-9: red; − residues position 45-49: blue).

Equivalently, it is hypothesized that the elongation proceeds at a faster rate when residues 5 to 32 are incorporated at the PTC, and is slower when residues 33 to 59 are incorporated at the PTC. The impact on the elongation speed will be quantitatively assessed with the use of a Maxwell-Boltzmann factor. This Maxwell-Boltzmann factor provides a quantitative modulation of the average elongation speed (see supplemental material).

It is also hypothesized that the ribosome requires more mechanical power to push the nascent chain out of the tunnel when it is repelled due to the electrostatic interactions, when residues 33 to 59, in our example, are incorporated at the PTC. How the extra mechanical power is mobilized is currently unknown. An increased turnover in the biochemical reactions providing Gibbs free energy to the ribosome would probably help. Equivalently, this would require an increased rate in amino acid incorporation because more Gibbs free energy would then be available as there are two energy rich GTPs hydrolyzed per residue incorporation.

Fig. 10 (B) shows the asymmetric potential (realistic model) effect on 5 contiguous positively charged residues between position 5 and 9 (net positive charge centered at position 7). The nascent peptide is attracted into the tunnel until amino acid residue number 19 (= 7+12) is incorporated at the PTC. From position 20 to 26 = 9+17, the axial forces acting on the peptide tend to pull it back into the tunnel and these forces tend to prevent the peptide from moving out of the tunnel. Then again, from position 27 to 37, the axial forces acting on the peptide tend to move it out of the tunnel. Finally, from position 38 to 58, the axial forces acting on the peptide tend to pull it back into the tunnel and these forces tend to prevent the peptide from traversing the tunnel easily. Compared with Fig. 10 (A), there are two fast moves separated by a short slower move, before residue 38, instead of one single fast move in the symmetric potential case. Equivalently, it is hypothesized that the elongation proceeds at a faster rate when residues 5 to 19 then 27 to 37 are incorporated at the PTC, and is slower when residues 20 to 26, then 38 to 58 are incorporated at the PTC. Note the amplitude of the axial forces are smaller but more dispersed in the positive region, for the asymmetric potential (realistic model) (B), than for the symmetric potential (A). Fig. 10 (C) and (D), similarly show the same effects but for 3 contiguous arginine residues instead of 5, (A) and (B). The amplitudes of the axial forces are 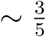 for (C) and (D) as compared to (A) and (B). Fig. 11 (A) shows the symmetric potential (idealized model) effect on a peptide with 5 contiguous positively charged residues between position 5 and 9 (net positive charge centered at position 7) and 5 contiguous negatively charged residues between position 45 and 49 (net negative charge centered at position 47) exactly 40 residues away from the first charge cluster. Fig. 11 (B) shows the asymmetric potential (realistic model) effect on the same peptide sequence. When the first plus cluster is emerging at the tunnel exit point, the second minus cluster is at the tunnel entry point. This situation results in high axial forces making difficult for the peptide to get out of the tunnel. As for Fig. 10 (B) and (D), there are two fast moves separated by a short slower move, before residue 38, instead of one single fast move in the symmetric potential case. The estimated maximal axial force is more than 8 pN and is reached when residue 52 is at the PTC. The axial forces tend to prevent the peptide to get out of the tunnel when residues 38 to 59 are at the PTC.

#### 2. Global and local electrostatical work and energy balance

In the two symmetric electrostatic models (cylinder and cylinder with Lorentzian peak), it is important to note that there is no difference in electrostatic potential between the entry point and the exit point of the ribosome exit tunnel, whatever the form of the potential inside of the tunnel. Because electrostatic interaction is conservative, the total work spent or harnessed by a charged residue when moved from the entry point to the exit point of the tunnel will always be equal to zero. Hence, the global net mechanical work for a full sequence to be moved completely through the ribosome exit tunnel should always be equal to zero. In the two asymmetric electrostatic models (cylinder plus truncated cone with or without the Lorentzian peak), there is a net difference in electrostatic potential between the entry point and the exit point of the ribosome exit tunnel. The total work spent or harnessed by a charged residue when moved from the entry point to the exit point of the tunnel will not be equal to zero in general. With a potential difference of 1.7 mV between the tunnel exit and entry points, the required mechanical energy is −0.164 kJ/mol (0.039 kcal/mol), or ~ 0.3 pN · nm on a single molecule, to traverse a single positively charged amino acid residue through the tunnel. Moreover, in any case, transiently or locally, the work to overcome positive axial electrostatic forces or the work harnessed in case of negative axial electrostatic forces acting upon any unit charged test residue may not be equal to zero. To illustrate this, the local mechanical work is computed in the case of the simulated synthetic peptide of Fig. 11 (B) with 5 contiguous arginines (+) and 5 contiguous glutamates (−), separated by 40 neutral residues. When the oligopeptide stretch ranging from residue 5 to 19 is incorporated, the sign of the work is positive (work = +0.67 kcal/mol), according to our adopted conventions in Fig. 2, meaning that the stretch is freely benefitting electrostatic energy to traverse the tunnel during the incorporation of those amino acid residues. On the contrary, when amino acid residues 38 to 59 are incorporated in the nascent chain, the sign for the work (work = −1.42 kcal/mol) is negative, meaning that mechanical energy has to be provided in some way to the nascent chain to help the stretch progressing through the tunnel. It is interesting to compare the computed values for the aforementioned mechanical work that are transiently either harnessed (0.67 kcal for the first stretch of 12 residues), or to be delivered (−1.42 kcal = 9.9 pN · nm for the second stretch of 21 residues), to the Gibbs free energy released from biochemical reactions at each residue incorporation, i.e. ΔG° ~ −18.3 kcal/mol (per amino acid incorporation) as detailed in the supplemental materials. If the chemical energy to mechanical work conversion yield is of the order of ~ 50%, an estimate of the local required chemical energy to push the nascent chain in the case of the second stretch would be around −1.42/0.5 = −2.84 kcal/mol. This amount of biochemical energy is about ~ 15% of the Gibbs free energy released from the biochemical reactions by a single new residue incorporation associated to the ribosome elongation cycle. These simple rough comparisons show that, energetically, the ribosome has enough energy resources to overcome the local electrostatic barrier easily.

However, situations may occur for which a nascent peptide will pose difficulties to the ribosome, considering that as much as ~ 15 %, or possibly more than ~ 30 % of the Gibbs free energy normally available to the ribosome per elongation cycle could be required to push the nascent chain out of the tunnel, depending on the charged amino acid distribution content of the nascent chain, and depending on the section widening in the region close to the exit point of the ribosomal tunnel.

#### 3. Real protein sequences

The purpose of the ribosome exit tunnel electrostatic realistic model is to apply it to real protein sequences, to compare them and to quantitatively determine where are the critical spots for the ribosome elongation process, or what are the axial force profiles acting on proteins during the co-translational folding process. To illustrate the application of our model to compute the forces acting on real protein sequences, we use it here in the context of neurodegenerative diseases like Huntington’s, Creutzfeldt-Jakob, or Alzheimer’s diseases. These diseases share a common pathology in the deposition of misfolded and aggregated conformations of a particular protein in the central nervous system at sites of neuronal degeneration [47]. The mechanisms of misfolding, aggregation and their functional consequences are not yet fully elucidated. Huntington’s disease is caused by mutations that expand the number of glutamine codons within an existing poly-glutamine (poly-Q) repeat sequence of the gene coding for the huntingtin protein [47–50]. The N-terminus end of a normal huntingtin protein is composed of a N-terminus sequence of 17 residues (NT_17_), a poly-Q sequence with a number of contiguous glutamines anywhere between 6 and 34 (e.g. Q_21_), and a polyproline sequence of around 11 proline residues (e.g. P_11_); see Fig. 12. A mutant allele coding for a number of glutamine repeats exceding 36 (e.g. Q_36_) will inevitably lead to Huntington’s disease if the person carrying this allele lives long enough. Huntingtin has a very long sequence with a total length of 3, 144 residues in the normal wild type sequence but the mutated huntingtin is only expanded in the very beginning of the sequence. Here, we do not pretend to solve the mechanism or the detailed molecular steps causing the misfolding of the huntingtin mutant protein but provide an analysis of a possible role of the forces acting on the huntingtin growing sequence while it is biosynthesized by the ribosome and investigate a possible co-translational misfolding situation. We compare, in Fig. 13 (A) and (B), the axial forces profiles for the human wild type huntingtin `HTT` and a mutant huntingtin `mHTT` for the first 150 N-terminus residues when their respective transcripts are being translated. The folding conditions and environments are different as the axial forces acted by the exit tunnel of the ribosome on these two growing nascent huntingtins are different. The two sequences embedded in the tunnel are different and cause the two very different net resulting axial forces. The mutant huntingtin has a length of the N-terminus sequence equal to 64 (= *NT*_17_ + *Q*_36_ + *P*_11_).

**FIG. 12:**
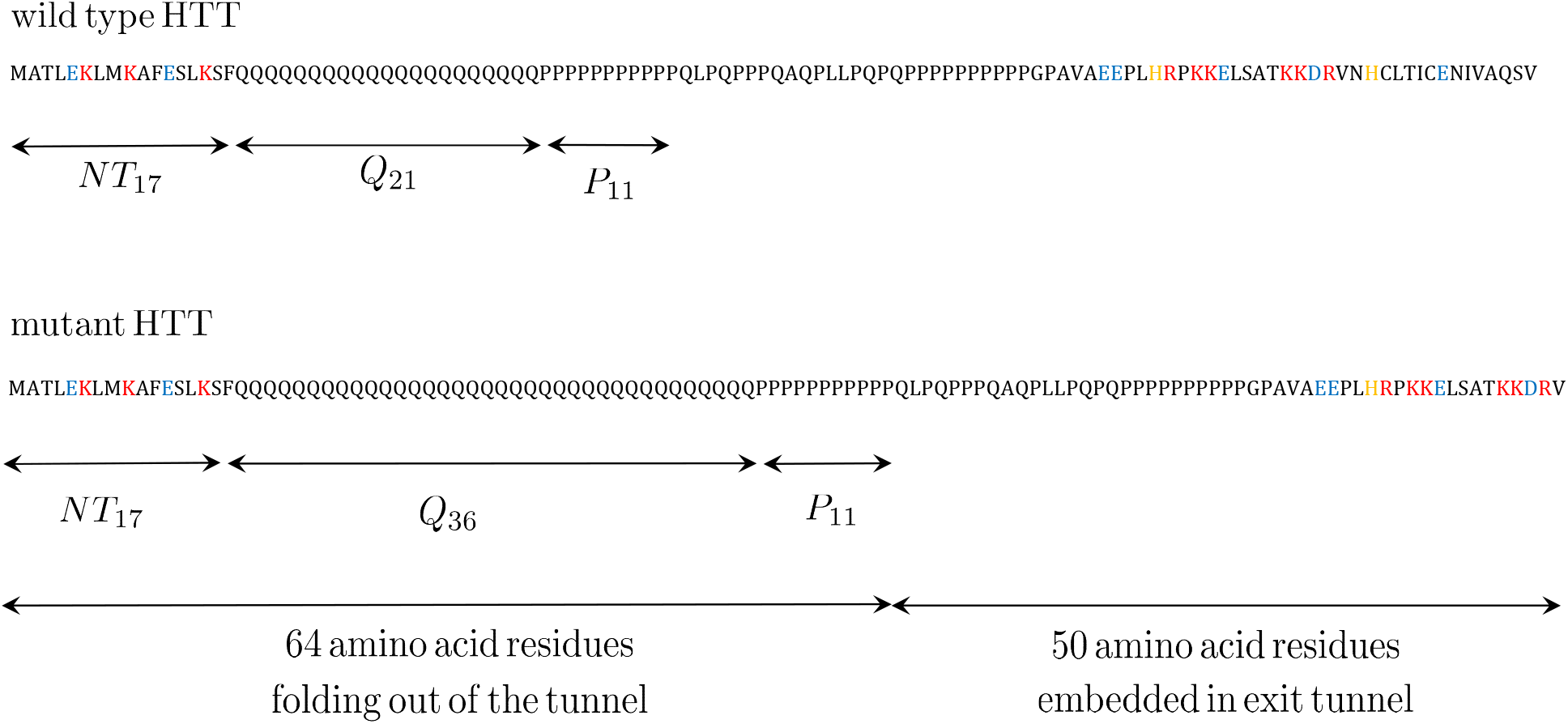
Wild type HTT and mutant `mHTT` huntingtin protein N-terminus starting sequence, showing the lengths of their poly-Q sequences. Positive residues: red. Negative residues: blue. Histidine residues: orange. Neutral residues: black

**FIG. 13:**
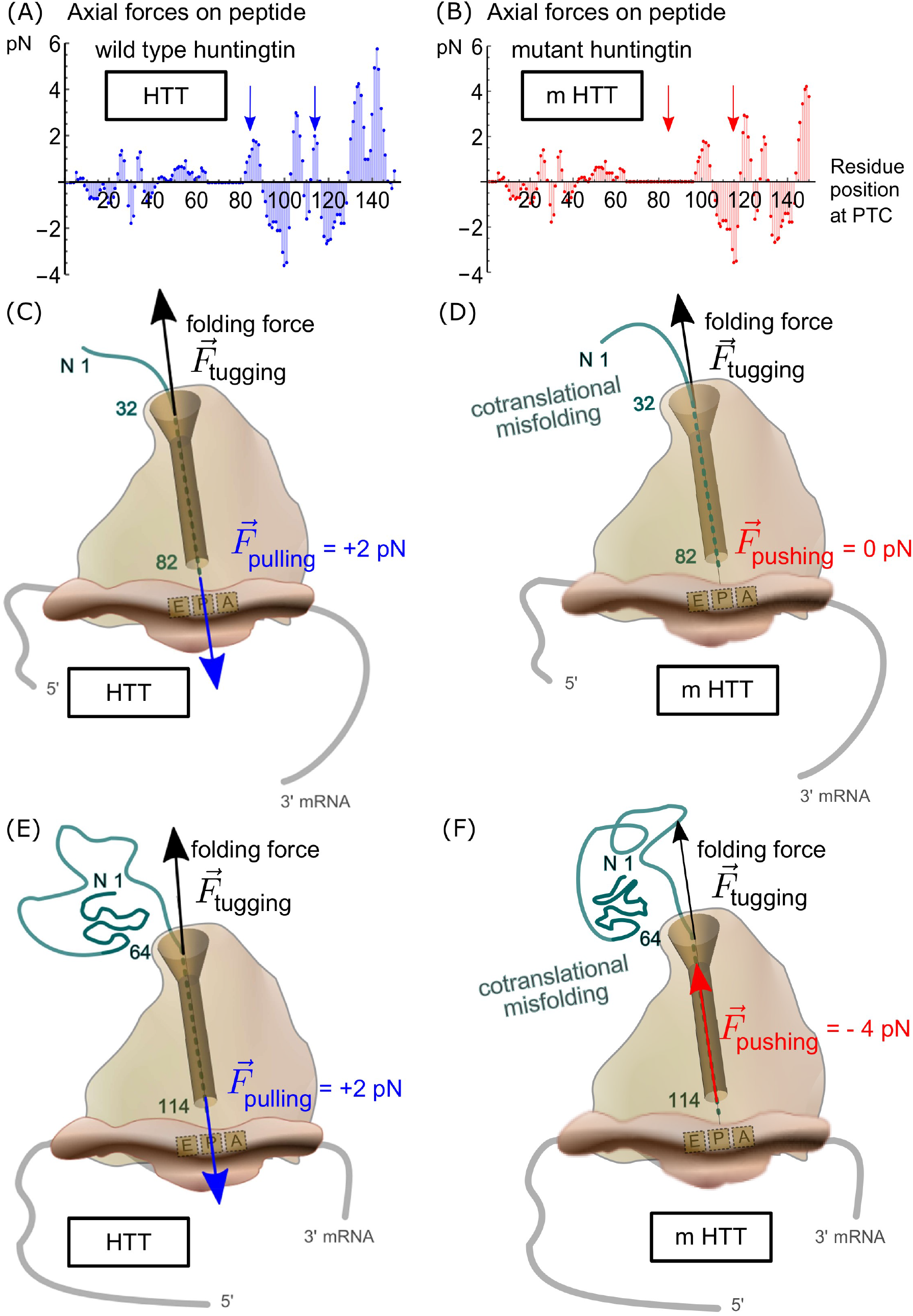
(A) and (B) Axial forces profiles for wild type and mutant protein. Blue and red arrows show the values of forces acting on the nascent chain at the PTC when residue 82 and 114 respectively are incorporated in the peptide at the PTC. (C) Wild type protein. A 2 pN pulling force due to the tunnel interaction with residues 32-82 opposed to the spontaneous folding force when residues 1-31 are out of the tunnel. (D) Mutant protein. No force opposed to the spontaneous folding force when residues 1-31 are out of the tunnel. (E) Wild type protein. A 2 pN pulling force due to the tunnel interaction with residues 64-114 opposed to the spontaneous folding force when residues 1-63 are out of the tunnel. (F) Mutant protein. A pushing force of 4 pN adds to the spontaneous folding force. Comparison of vectorial co-translational folding for huntingtin wild type (HTT) and mutant protein (mHTT).

The exit tunnel exerts axial forces of electrostatic origin that are either pulling forces or pushing forces. These forces oppose (or not) to the forces generated by the spontaneous folding of the unstructured segments of the nascent polypeptide chain upon lengthening of the chain out of the tunnel. In their computational simulations, Fritch *et al.* estimated that the force difference experienced at the P-site residue upon doubling the length of the chain out the tunnel was of the order of piconewtons [51]. In our Fig. 13 (C) to (F), these folding forces are called tugging forces (black arrows). In the extreme case of SecM mediated ribosomal arrest known in bacteria, Goldman *et al.* provided evidence that the minimal tugging force must be ~ 10 pN to relieve the stalled ribosome [52]. These results show that tugging forces in the range 1 − 10 pN can possibly be generated by emerging folding nascent chains to reach their native conformation.

As seen from the comparison of the axial forces profiles, the wild type nascent protein is experiencing a pulling force (~ +2pN) of electrostatic origin from the ribosome, when incorporating residue 81 to 90 at the peptidyl transferase center PTC, Fig. 13 (A) first arrow, and Fig. 13 (C), while the length of the nascent chain out of the tunnel is 31 to 40, i.e. when the critical poly-Q segment is fully emerging from the ribosome. In the mutant `mHTT`, there are no pulling forces from the ribosome at this moment; the axial forces are null at this moment, Fig. 13 (B) first arrow, and Fig. 13 (D).

When the first 64 amino-terminal residues have just emerged out of the exit tunnel of the ribosome and are exploring the folding space, the downstream 50 residues are under electrostatic interaction with the ribosome exit tunnel. The total axial resulting force acting upon residue 114 (= 64 + 50), and indirectly on the whole growing chain outside the tunnel, is different for the mutant `mHTT` than for the wild type `HTT`. For the wild type `HTT`, the nascent chain out of the tunnel, 64 residues in length, folds while the ribosome is pulling the chain toward the interior of the tunnel (+2 pN), Fig. 13 (A) second arrow, and Fig. 13 (E), whereas for the mutant protein `mHTT`, the nascent chain out of the tunnel, 64 residues in length, folds without opposing force from the ribosome. On the contrary, for the latter, there is a pushing force on the nascent chain (−4 pN) from the ribosome exit tunnel, Fig. 13 (B) second arrow, and Fig. 13 (F). This analysis would suggest that proper folding of the poly-Q containing segment of huntingtin protein would require a pulling force from the ribosome. If there is no pulling force, as for the mutant huntingtin, the poly-Q segment would be much more prone to co-translational misfolding. Interestingly, the effect of expanding the length within the poly-Q segment would just cause a shift in the axial forces profile that reverse the forces acted by the ribosome, upon the folding segment, between the wild type and the mutant protein.

Overall, these results suggest that a change in the local distribution of charged residues or an insertion or a replacement by neutral residues has impact on the axial forces profile over a spatially extended region of the nascent protein which is in a range corresponding to the length of the ribosome exit tunnel. The proper co-translational folding of a nascent polypeptide chain out of the ribosome calls for specific patterns in the charged amino acid distribution in the sequence downstream, embedded in the exit tunnel, down to the peptidyl transferase center PTC. The encrypted sequence indirectly dictates, in a spatially extended way, the electrostatic interaction of the charged amino acid residues in the exit tunnel to generate axial forces profiles acted by the ribosome upon the growing chain. These forces would play a key role in the correct co-translational folding process. The co-translational folding is vectorial, that is, it involves elements that emerge successively from the N-terminus to the C-terminus [53]. The landscape of co-translational folding may differ depending on the charged residue distribution which is embedded in the tunnel. Our model sheds light on how the ribosome could affect the folding trajectory.

## V. DISCUSSIONS AND FUTURE PERSPECTIVES

In this study we set out to model the electrostatics of the ribosome exit tunnel to explore quantitatively the impact of the distribution of the charged amino acid residues embedded in the tunnel on the forces acting on the nascent peptide chain during translation. Our approach was to develop a full analytical expression of the electrostatic potential inside the tunnel, starting from two idealized theoretical geometries for the tunnel, i.e. a cylinder and a cone. We eventually concatenated the cylindrical geometry with the conical geometry, and finally added an empirical Lorentzian function motivated by the known experimental observations of local and highly conserved ribosomal protein protrusions inside the tunnel. The precise geometry of the tunnel is important for quantifying the resulting electrostatic potential profile. It shows what part of the electrostatic profile is contributed by the shapes and by the sizes of the tunnel and what part is inherent to the physicochemical properties such as the surface charge density contributed by the large number of phosphates moieties lining the tunnel inner wall.

The main results derived from the theoretical analysis of the electrostatics of the ribosome exit tunnel, as displayed in Fig. 7, Fig. 8 and in Fig. S1 (supplemental material), is the goodness of the fit of the physical model with the measured data points for the electrostatic potential in the ribosome exit tunnel earlier published by Carol Deutsch and co-workers [29]. The geometry and physicochemical properties of the tunnel inner walls of the exit tunnel consistently explain the experimentally measured values for the potential from simple first physical principles. The model draws the attention on the main geometrical and physical features as determinants of the electrostatic potential profile and the derived electric field projected along the tunnel axis. Specifically, the geometrical variation induced by widening the tunnel radius at the exit of the tunnel (cone frustum) introduces a permanent difference in the electrostatic potential between the exit and the entry points of the tunnel. This is energetically unfavorable to the positively charged amino acid residues as compared to their negatively charged amino acid counterparts. This provides a simple bioenergetic explanation to the observation that, proteome wide and across species, the protein sequences are slightly but significantly more enriched in negatively charged amino acid as compared to the positively charged amino acid residues [54]. This observation would be the consequence of a selection pressure in favor of the negatively charged residues as compared to the positively charged residues; the latter requiring more mechanical energy to traverse the ribosome exit tunnel. The derived axial forces acting upon the nascent polypeptide stretch, within 50 residues upstream of the amino acid residue that is incorporated at the PTC, stand as a valuable quantitative model. The variation in the axial forces due to electrostatic interaction of the charged nascent chain with the ribosome exit tunnel has been estimated in a range from −10 pN to +10 pN in order of magnitude. More importantly, different profiles for these axial forces have been quantitatively related to synthetic polypeptides with arbitrarily charged residue distribution. Arbitrary synthetically engineered transcripts could, in principle, be used in high resolution optical tweezers multiple traps experiments to test experimentally the theoretical profiles of the axial forces acting upon such nascent polypeptides. The electrostatic model best fitted to the experimental data of Lu *et al.* [29]. is the one combining a cone frustum section concatenated to a cylindrical section with a Lorentzian peak roughly located one third of the tunnel length away from the tunnel entry point. This particular model is used to derive, more accurately, the axial forces acting upon any nascent chain in the tunnel. The comparison of the axial force profiles of wild type protein sequences with mutant sequences as illustrated in the case of huntingtin (Fig. 12 and Fig. 13) may help to study the dynamical folding of a nascent protein that is still in contact with the ribosome. The tugging forces generated by the spontaneous folding of the unstructured segments during the peptide lengthening out of the tunnel were estimated by computer simulations at piconewtons order of magnitude by Fritch *et al.* [51]. These spontaneous co-translational folding tugging forces acting on the nascent chain can be compensated for (or not) by pulling forces from the ribosome due to the electrostatic interaction in the ribosome exit tunnel. The landscape of co-translational folding of the wild type and mutant huntingtin nascent proteins may differ by a sheer difference in the distribution of the charged amino acid residues that are embedded in the full length of the tunnel. This would shed light on how the ribosome takes part in configuring folding intermediates [53]. The specific pattern of the axial forces acting on the residues that are incorporated successively from the N-terminus to the C-terminus could prevent the emerging nascent chain from falling in kinetic traps, or in stable misfolded conformations, eventually resulting in protein aggregation. Our model allows a quantitative analysis of these axial forces profiles and a comparison of such profiles between correctly folded and misfolded protein conformations. The ordered list of axial forces at single residue resolution also allows to calculate the mechanical work required to overcome the electrostatic potential real profile in the exit tunnel at each residue elongation. From this, a Maxwell-Boltzmann correcting factor can be defined following similar developments as the ones exposed in [55, 56] and introduced in a seminal article by Bell in the context of cell to cell adhesion [57]. These factors can correct, at single amino acid residue resolution, and in a sequence specific way, the elongation rate in TASEP-like modeling tools. For a given transcript, the specific contribution of the tunnel electrostatic interaction locally modulates the elongation rate in a range from minus 40% to plus 85% when compared to the average elongation rate; see supplemental material (C). An interesting advantage of these Maxwell-Boltzmann correcting factors lies with the way they are calculated. The exact local memory of the distribution of the charged amino acid residues is conserved for a sliding window of 50 residues that are upstream the site of incorporation of a new residue at the peptidyl transferase center PTC. This extended stretch of 50 residues is expected to be under the influence of the electrostatic interaction caused by the inner wall of the ribosome exit tunnel, along its whole axial length. All charged residues, positive and negative, embedded in the tunnel, additively contribute to the pace of the elongation process. The route of force transmission to the P site residue is through the nascent polypeptide’s backbone as it is also the case for the tugging force generated by the spontaneous folding of the lengthening nascent chain out of the ribosome exit tunnel [51].

Mechanical forces can alter the activation energy barriers that reactants have to overcome in the course of a chemical reaction to be converted into products. Intermediate transition states may be more easily attainable from the reactants when the system is experiencing an external force [55]. An effect of the external applied force is to provide mechanical work that will linearly decrease the activation energy even without changing the reactants’ configurations or the transition state configuration [56, 57]. When the axial forces upon the nascent chain buried in the tunnel are exerted toward the tunnel exit, the Gibbs free energy barrier at the PTC is presumed to be decreased, the rate of the peptidyl-tRNA deacylation step at the P site and the global rate of the peptide bond formation are both expected to be increased.

To our knowledge, the model presented here is the first one to take into account the whole size and shape of the ribosome exit tunnel and updates, at single residue resolution, the mobile 50−mer polypeptide window which is embedded in the tunnel. The position dependent precise value of the Maxwell-Boltzmann factor is determined by this spatially extended stretch of 50 residues with a specific charges distribution that is encrypted in the transcript being decoded. These elongation rate correcting factors are at codon resolution and keep the memory of the spatially extended stretch of amino acid residues embedded in the tunnel. This is a clear improvement over the current state of the art in terms of realism and consistency of the elongation speed calculation. This can be contrasted, for instance, with studies where only positively charged residues within a limited number of residues upstream the incorporation site are considered and where arbitrarily fixed valued correcting factors are used to adjust for the electrostatic interaction in the tunnel [14].

Our model of the electrostatic interaction of the ribosome exit tunnel with the nascent chain polypeptide relies on a number of critical assumptions which prevent to consider the model as a completely realistic representation. As advocated by Lucent *et al.* [40], the understanding of the complexity of molecular behavior in the ribosome exit tunnel should require an atomistic molecular dynamical description including the solvent confined to the tunnel as the medium inside the tunnel does not behave as a continuous isotropic dielectric medium. In our simplified approach, we nevertheless considered the isotropic dielectric permittivity of the tunnel medium to be the one of water as a bulk-like solvent (*ϵ* = *ϵ*_0_ · *ϵ*_*r*_ where *ϵ*_*r*_ = 74 at 37 degrees Celsius). The exact size of the exit tunnel and the number of accommodated residues inside the tunnel are not known with full certainty, especially in mammals. We relied only on the pioneering works made on prokaryotes by Voss *et al.* [28] and Lu *et al.* [29]. Of course, the model could still be adjusted according to new experimental evidence regarding the exact number of residues embedded in the tunnel or to improved resolution in the electrostatic potential profile measurement inside the tunnel. The possible secondary structures that can start to form near the exit end of the tunnel have not been taken into account in our study. Existing X-ray crystallography and cryo-electron microscopy data should be used more intensively to corroborate the values of our model main parameters both for the size and net charge surface density. Furthermore, the available electron microscopy data should challenge the uniformity in the surface charge density on the tunnel inner wall that we assumed and the possible differences between the surface charge densities of the cone frustum inner wall section, *σ*_2_, and the cylindrical inner wall section, *σ*_1_. It is also not known with full certainty whether or not the shape and geometry of the ribosome exit tunnel in the large subunit LSU of the ribosome stay the same during the translation process in vivo or if reversible continuous elastic deformations occur in vivo. We specifically showed that a dynamical change in the opening angle of the cone frustum at the exit tunnel would result in a dynamical variation of the electrostatic potential at the axial exit point of the tunnel. The electrostatic potential profile is sensitive to the tunnel geometry. The molecular basis of these hypothetical dynamical conformational changes in the exit tunnel shape is still to be elucidated. Auxiliary chaperone proteins could assist the large ribosome subunit LSU to change the opening angle of the cone frustum. An intrinsic chemical modification of a LSU component itself could alter the shape of the exit tunnel. These events could be elicited during ribosome stalling rescue mechanisms. Most likely, the hypothetic reversible elastic deformation in the cone frustum angle at the tunnel exit could be concomitant to the internal reversible conformational changes in the large ribosome subunit LSU during the transpeptidation reactions occurring in the LSU and induced synchronously when the ternary complex (aa − tRNA • EF • GTP) is accommodated at the A site and eventually at the release of the elongation factor, the guanosine diphosphate (GDP) and the inorganic phosphate. We showed that the mechanical energy required to push the growing nascent chain through the LSU exit tunnel, even in difficult scenarios, would be smaller than the Gibbs free energy released from the transpeptidation and the hydrolysis of a single GTP. Overall, the widening of the radius along its central axis toward the exit of the tunnel is however known and contributes to the asymmetric electrostatic potential profile that we estimated. This estimated electrostatic potential profile fits the available observed data rather well at least for prokaryotes. We must recognize, however, that we only relied on a small sample size of 4 to 6 point measurements. Complementary wider experimental studies on both prokaryotes and eukaryotes ribosomes would be beneficial.

The electrostatic potential mathematical model that we proposed provides insights into the real measurements that were made in the pioneering experimental studies and that could still be made in the future. It should be emphasized that electrostatic potential measurements should always be conducted in association with precise measurements of size and shape of the tunnel and accurate positional mapping along the tunnel axis. To quantitatively estimate the axial forces applied on the nascent chain, we made a rigid body assumption or assumed the non-deformability of the nascent chain inside the tunnel. This assumption is most certainly not valid for all polypeptide chains and most probably not valid locally. However, this assumption could be legitimate on average and proteome wide. Indeed, the ribosome exit tunnel is universal, meaning that all the polypeptides that are naturally occurring in the biosphere did traverse the tunnel at the time of their biosynthesis. All the amino acids have progressed through the entire length of the tunnel after they were incorporated in the nascent chain at the peptidyl transferase site. On average, as a first approximation, we can consider that these amino acids followed a centro-axial trajectory in the tunnel and experienced the effect of the electrostatic interaction upon the charged residues with which they are directly or indirectly bound. Fritch *et al.* recently showed that the spontaneous folding force was transmitted directly from the outside of the tunnel to the PTC center through the backbone of the nascent chain [51]. This direct transmission route supports the rigid body assumption we made for the peptide buried in the tunnel. On average, it is believed that the comparison of the electrostatic interaction with the exit tunnel of any two different nascent chains, by applying our model, can provide quantitative insights on the effects of the difference in charged amino acid distribution across their primary sequences. This paves the way to a variety of bioinformatic studies on transcriptomic and proteomic data to shed light on translational control. We expect the in-silico research community to assign itself the task of using our suggested electrostatic model, and the ordered list of Maxwell-Boltzmann factors derived from it, to modulate the elongation rate for a better quantitative account of the effect of the tunnel on the charged amino acid residues. Immediate perspectives and objectives will address (a) accurate predictions of ribosome footprints in Ribo-Seq profiling ensemble experiments; (b) precise dynamical predictions in the speed of elongation in single mRNA molecules experiments; (c) quantitative predictions of the measured tugging force profiles on nascent polypeptide chain emerging from the ribosome exit tunnel in high resolution multiple traps optical tweezers experiments to be conducted on tethered ribosomes in vitro; (d) comparison of axial forces profiles associated to correctly folded or misfolded proteins for the study of co-translational folding and protein aggregation mechanisms. The model presented in this study consistently connects different results and experimental observations coming from different fields in molecular biology, structural and physical chemistry, synthetic and multi-omics biology and provides a clear picture of the electrostatic interactions in the ribosome exit tunnel and their effects on the protein elongation rate.

## VI. SUPPLEMENTAL MATERIALS

### A. Normally truncated straight cone model complete derivation

#### 1. Scalar potential

The scalar potential for the normally truncated straight cone model was expressed by equation (34) as

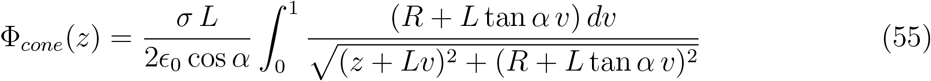

The expression inside the square root in the denominator of the integrand can be written

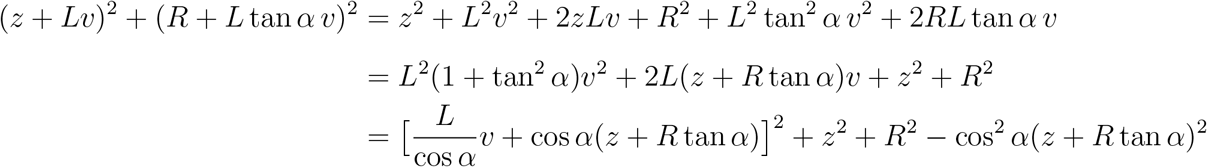

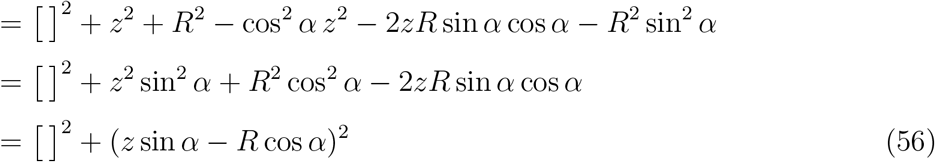

and so the square root in the above denominator can be rewritten

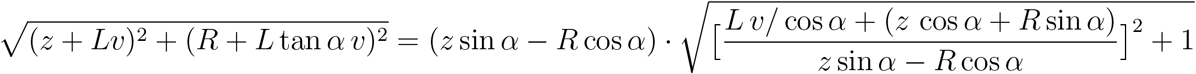

To alleviate the notations, we pose as in (35) and (36)

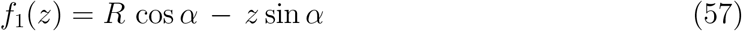

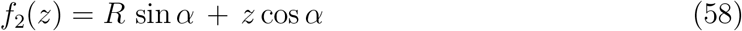

and we pose

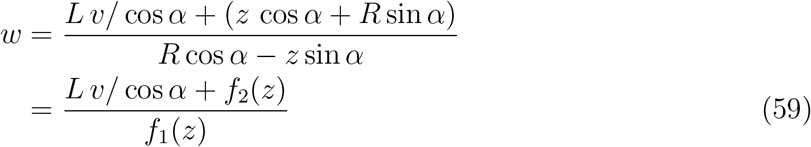

Hence,

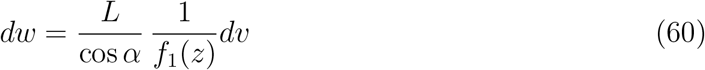

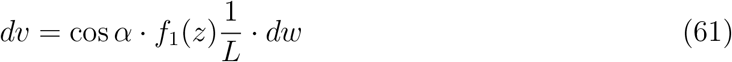

The numerator in the integrand of (38) now writes

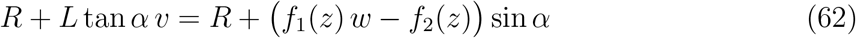

and equation (34) turns into

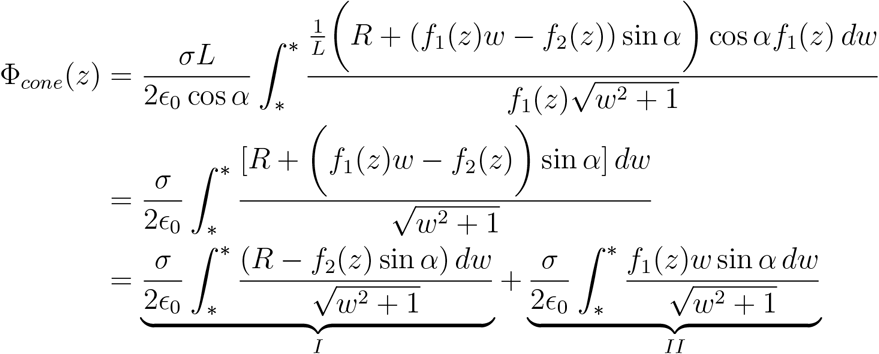

These two terms easily integrates. The first one (I) is still simplified further through

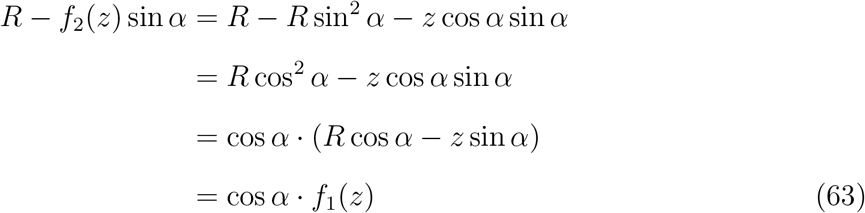

and so,

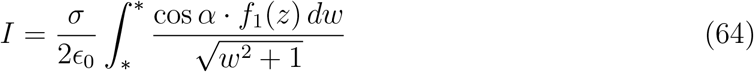

Substituting *w* = sinh *y*, *w*^2^ + 1 = cosh^2^ *y* and *dw* = cosh *y dy*, yields

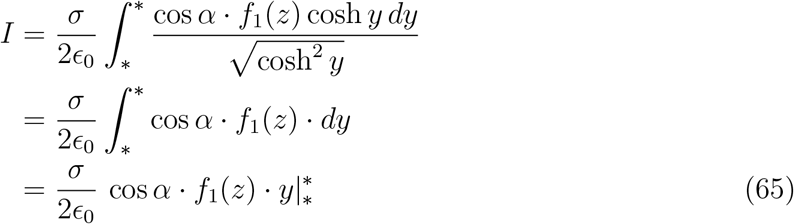

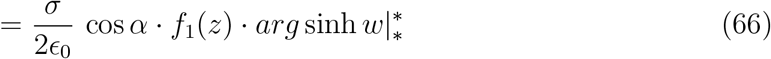

but 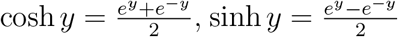 and cosh *y*+sinh *y* = *e^y^*, so 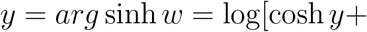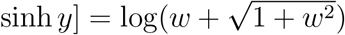. Hence,

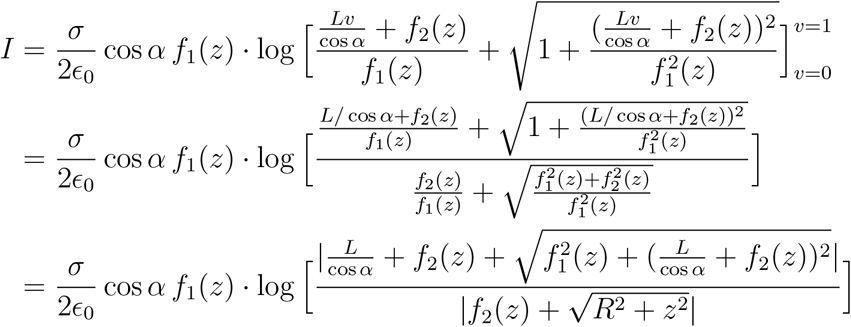

Noticing that

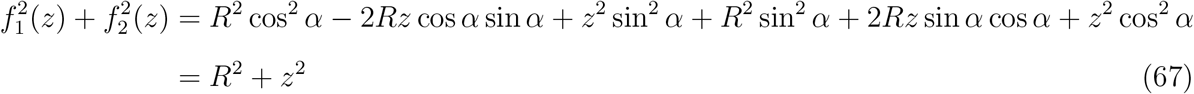

and that

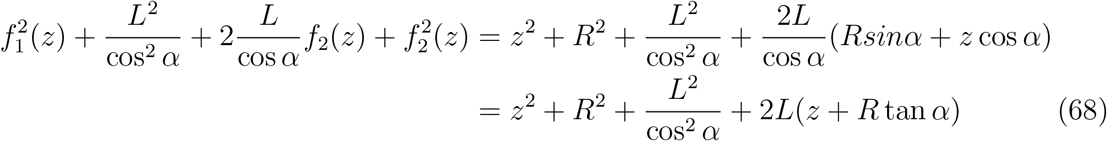

results in

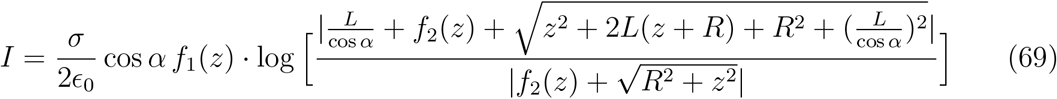

For the second term (*II*), substituting *w* = sinh *y*, cosh^2^ *y* = sinh^2^ +1 = *w*^2^ + 1 and *dw* = cosh *y dy*, we have

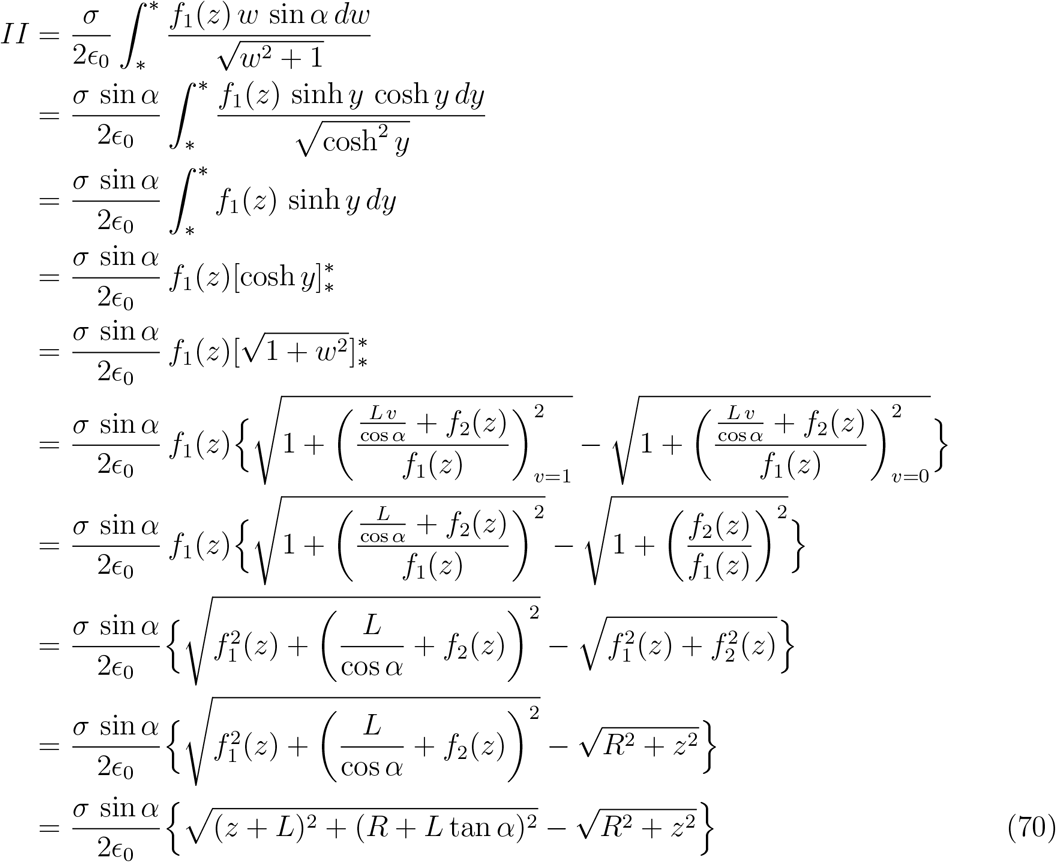

Summing the two terms I and II results in equation (38).

#### 2. Electric field

The complete derivation of the electric field projected along the tunnel axis follows from

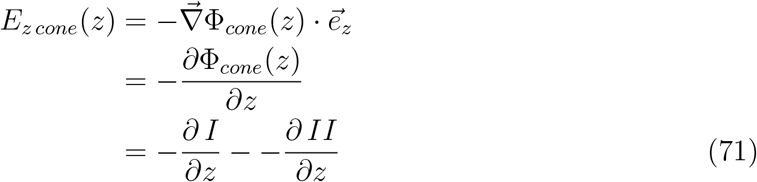

We start with 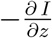

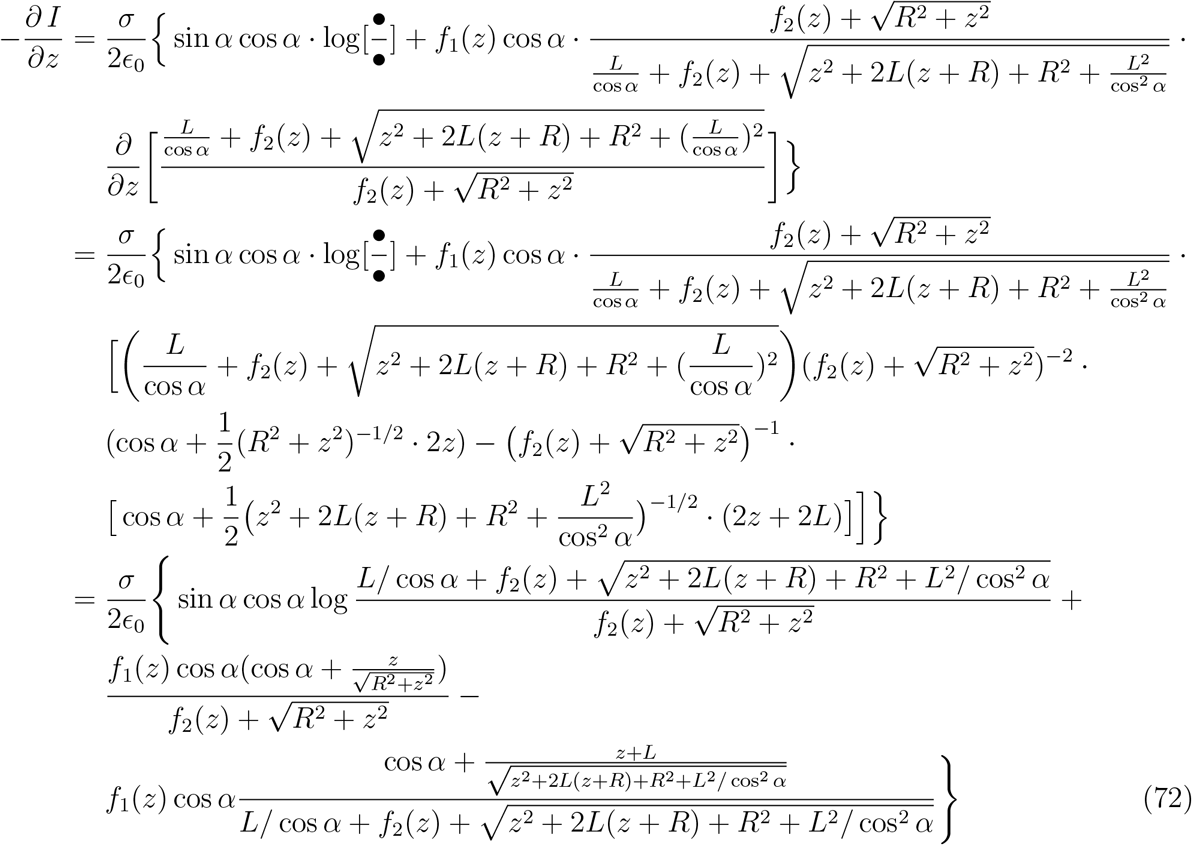

We go on with 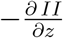

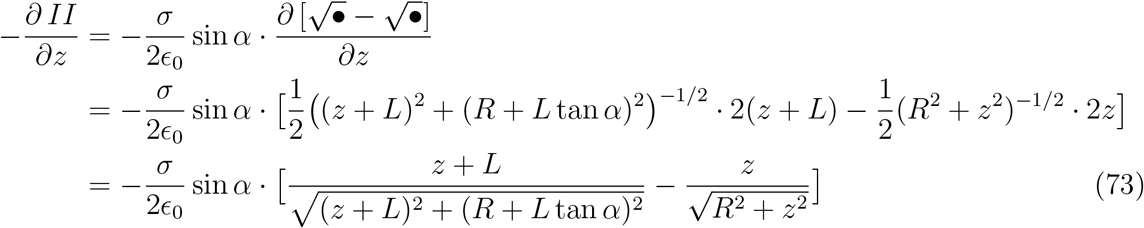

Summing the two terms yields the final result for *E_z cone_*(*z*) as in equation (41)

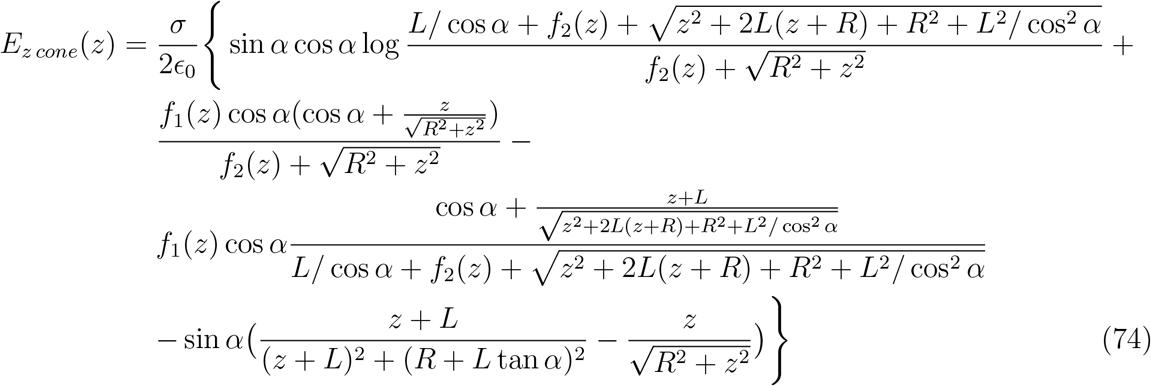

### B. Figure S1: animated figure showing dynamical changes in the exit tunnel cone frustum geometry

Figure S1, in supplemental material, shows how a dynamical change in the geometry of the opening angle in the cone frustum at the exit of the tunnel could change the electrostatic potential profile inside the tunnel. The animated figure is for a ribosome exit tunnel of length 8.5 nm. The animated figure Fig. S1 shows that an increase in the opening angle increases the potential difference between the exit and entry point of the tunnel. In this animated figure, the surface charge density of the cone frustum comply with the charge conservation and with the assumption of an elastic deformation of a preexisting cylinder surface turned into a truncated cone surface preserving the same total charge, see equations (52).

### C. Specific effect of the tunnel electrostatic interaction on the elongation rate

The electromechanical force due to the tunnel electrostatics acts on the peptide nascent chain and is transmitted inside the ribosomal tunnel up to the peptidyl transfer center (PTC) responsible for the peptide bond formation. The force is transmitted to the PTC through the whole length of the polypeptide chain backbone embedded in the tunnel [51]. At the PTC, the first event that must occur before the peptide bond is built between the peptidyl-tRNA at the P site and the aminoacylated tRNA at the A site is the breaking of the ester covalent bond between the oxygen atom attached on the tRNA 3’ end (3’ carbon at the CCA terminal ribose) and the carbonyl group of the carboxyl terminal end of the peptide. We presume that a force acting on the peptidyl-tRNA peptide directed from the P site toward the N-terminal end of the peptide would help breaking this ester bond. The chemical reaction rate of this ester bond breaking would be increased in the presence of such a force directed toward the exit tunnel. The ribosome elongation average rate can be quantitatively modulated by applying a Maxwell-Boltzmann factor, i.e 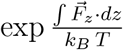, resulting from the theoretical treatment of the effect of force on the thermodynamics and kinetics of chemical reactions [55] or as initially introduced by Bell in a cell to cell adhesion context [56, 57]. This factor correcting the elongation rate specifically accounts for the electrostatic interaction of the nascent chain in the ribosome exit tunnel. This factor is calculated on the basis of all the 50 residues upstream and is updated at each new incorporation. The numerical value of this factor will be different at each residue incorporation and will always be dependent on the particular amino acid sequence being embedded in the tunnel. For the arbitrarily chosen protein `KIF4A` (member of the family of human kinesins), all the numerical values for the Maxwell-Boltzmann factors calculated for each residue sequentially incorporated at the PTC are displayed in Fig. 14. The minimal value is 0.60 and the maximal value is 1.85 for the Maxwell-Boltzmann factor in the particular case of `KIF4A`. 84.5% of the values are in the range [0.80, 1.20]. The mean, 1.01, is very close to 1.0. The minimal value of the Maxwell-Boltzmann factor, 0.60, occurs at incorporation of residue 1064 which is a E (negatively charged glutamate), in `KIF4A`, when the axial force on the nascent polypeptide stretch in the tunnel is +8.62 pN. The elongation rate is quantitatively slowed down by a factor 0.60. Equivalently, the time spent by the ribosome on codon 1064 is expected to be larger (average time for this type of codon divided by 0.60) at this position because of the most unfavorable electrostatic interaction occurring at the moment of this residue incorporation. The maximal value of the Maxwell-Boltzmann factor, 1.85, occurs at incorporation of residue 1103 which is a K (positively charged lysine), in `KIF4A`, when the axial force on the nascent polypeptide stretch in the tunnel is −10.49 pN. The elongation rate is quantitatively faster by a factor 1.85. Equivalently, the time spent by the ribosome on codon 1103 is expected to be smaller (average time for this type of codon divided by 1.85) at this position because of the most favorable electrostatic interaction occurring at the moment of this residue incorporation. This illustrates how the Maxwell-Boltzmann factors provide a consistent methodological tool to assess quantitatively the contribution to the elongation rate specifically due to the electrostatic interaction occurring in the tunnel, and in a separate way from the other factors affecting the mRNA translation rate.

**FIG. 14:**
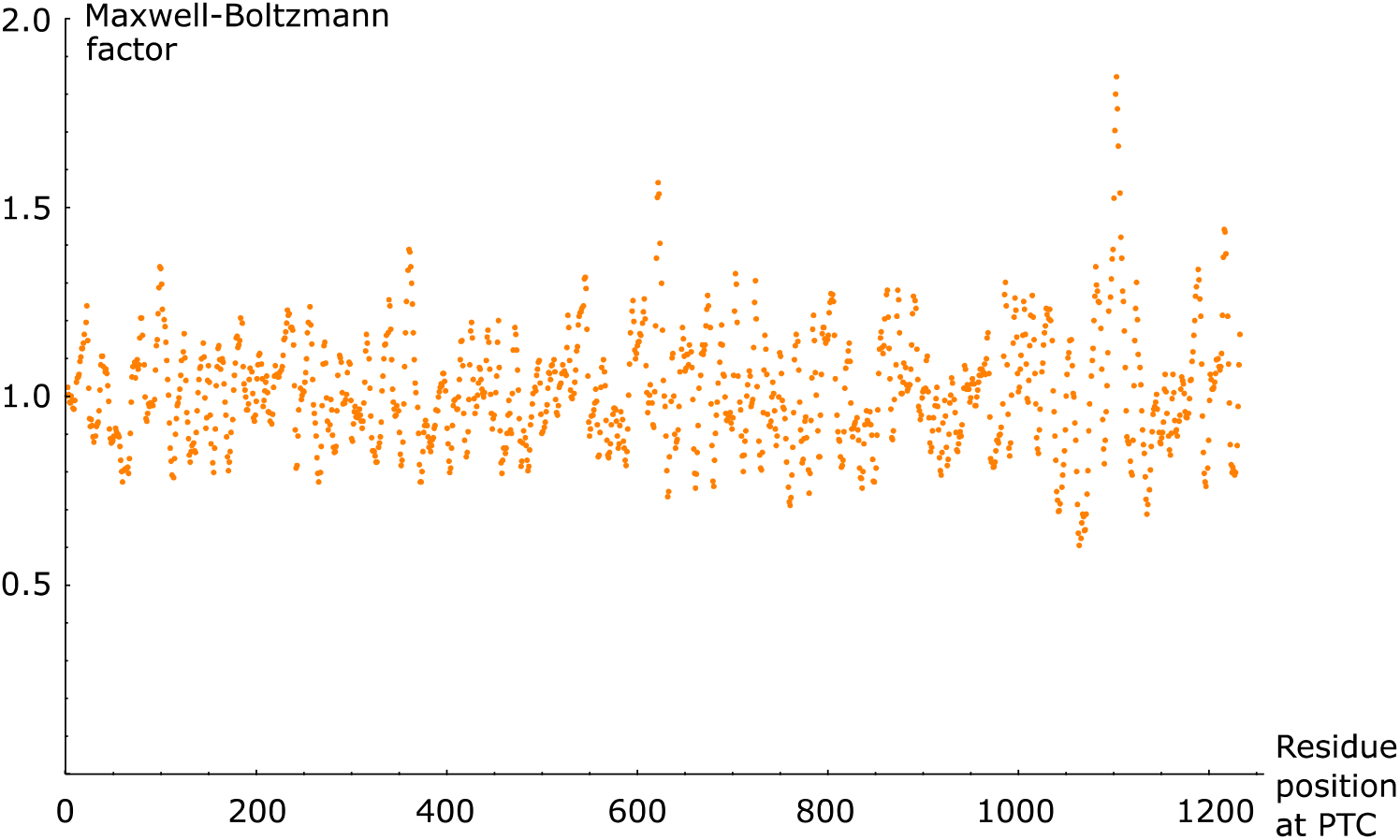
Maxwell-Boltzmann elongation rate factors weighting for the electrostatic interaction at each incorporation of a new residue at PTC as a function of residue position in the human protein KIF4A. Min = 0.60 and Max = 1.85 occurring at incorporation of residue *E*1064 and *K*1103 respectively. Lower values are associated to smaller elongation rate (slowdown), larger values are associated to higher elongation rates (speeding up).

### D. Energy sources available for the ribosome

The ribosome is a complex macromolecular machine that requires energy to carry out its multiple tasks. During elongation, a ribosome has to translocate the mRNA each time a codon has been paired to its cognate or semi-cognate tRNA and has to push the nascent protein through the exit tunnel.

The detailed energy balance (energy sources and uptakes) required for elongation has not been fully resolved. Our electrostatic model shows that, in certain situations, a Gibbs free energy fraction in the range 15% − 30% of the total biochemical energy available to the ribosome could be required to move the nascent protein through the exit tunnel.

The energy is found in the biochemical reactions taking place in the ribosome with the help of the associated catalytic sites of enzymes like the elongation factors (eEF in eukaryotes) or ribozymes. The elongation factors (EF and EF-G) are GTPases whose activity is controlled by the ribosome. When an aminoacyl group is hydrolyzed from the loaded tRNA, an ester group is broken and energy is released. For each amino acid incorporation cycle, two GTPs molecules are hydrolysed (one with the help of EF in the ternary complex accommodated at the A-site and one with the help of EF-G required for the mechanical translocation). The peptide bond formation itself requires free energy at each chain elongation by one residue. A very rough estimate of the net change in Gibbs free energy for the net balance between peptide bond formation and ester hydrolysis at pH = 7, 25°C yields ΔG° = −3.7 ± 1.2 kcal/mol = −15.5 ± 5.0 kJ/mol [32, 33]. This is known as the transpeptidation Gibbs free energy.

**Peptide bond formation:** the formation of the simplest dipeptide glycylglycine is endergonic and requires 15 kJ/mol (3.6 kcal/mol) per mole of formed peptidic bond: ΔG° = +3.6 kcal/mol for one residue incorporation (per ribosome cycle).

**Hydrolysis of ester bond in aminoacyl-tRNA:** the hydrolysis of the ester bond in aa-tRNA is exergonic and releases 30.5kJ/mol(7.3, kcal/mol) per amino acid released from the tRNA: ΔG° = −7.3 kcal/mol (per ribosome cycle)

**Hydrolysis of 2 GTPs:** the hydrolysis of 2 GTPs is exergonic and releases 30.5 kJ/mol (7.3 kcal/mol) per mole of GTP. Hence, per residue incorporation cycle (2 GTPs): ΔG° = −14.6 kcal/mole (per ribosome cycle)

**<u>Net Gibbs free energy available to the ribosome per aa residue incorporation</u>:** ΔG° = −18.3 kcal/mol (per residue incorporation)

The net result is that one ester bond to the 3’-hydroxyl of a ribose has been broken (locally in the ribosome) and one peptide bond in the nascent protein has been formed, two GTPs have been hydrolyzed, the ribosome has shifted forward the `mRNA` by one codon (translocation distance on `mRNA`, Δx ~ 1.4 nm (0.9 − 1.8), parenthesis indicate 95% confidence limits [33]) and the nascent peptide has advanced in the ribosome exit tunnel by one residue (nascent peptide chain distance displacement in the tunnel at each translocation, Δz ~ 0.25 nm, which is the estimated distance between two consecutive amino acid *α*−carbons as considered in our model). It is not fully elucidated whether (or how) free energy could be stored in the ribosome and used later to catalyze translocation and possibly assist the progression of the nascent protein through the ribosome exit tunnel when needed. Each step in translation involves intra-subunit or inter-subunit conformational changes [32–34]. Such conformational changes could store energy that could be released at a subsequent step, with a thermodynamical yield, providing a conceivable mechanism of harnessing the biochemical energy to use it for mechanical translocation and for moving the nascent peptide through the ribosome exit tunnel when required. The entropy driven spontaneous or chaperones assisted folding of the protein, generating a tugging force [22] outside of the ribosome exit tunnel, might also help the nascent protein to be pulled out of the tunnel. Optical tweezers assays have opened the way to characterizing the ribosome’s full mechanochemical cycle [33, 34]. Recently, such in vitro assays [33, 34] provided an estimate for the maximal mechanical energy required per translocation step (near stalling on the `mRNA`), 21.2 pN · nm = 5.2k_B_T, at 296 *K*, or ~ 3.1 kcal/mol. As estimated above, the Gibbs free energy available from the transpeptidation step (ester hydrolysis and peptide formation without the help of GTP hydrolysis) is ΔG° = −3.7 ± 1.2 kcal/mol. The mechanical work for translocation would be around 80% of the Gibbs free energy available from the transpeptidation. Such a high thermodynamic efficiency for conversion of chemical energy to mechanical motion is higher than occurs in most molecular motor [55]. Instead, efficient translocation would require the hydrolysis of at least one GTP with the help of elongation factor EF-G [33]. EF-G dependent GTP hydrolysis was shown to precede and greatly accelerate translocation [58]. The mechanical translocation of the ribosome on the mRNA by one codon would take 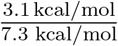 or 43 % of the Gibbs free energy released by the hydrolysis of one GTP, assisted by elongation factor EF-G. The mechanical energy required to push the nascent peptide chain through the large subunit exit tunnel could be provided by the transpeptidation Gibbs free energy or the hydrolysis of one GTP assisted by elongation factor EF in the ternary complex accommodated in the A site or a combination of both.

## Supporting information

Supplemental Figure 1: animated figure of ribosome exit tunnel

## Acknowledgments

We gratefully acknowledge Professor John Martin for fruitful discussions and suggestions on the electrostatic idealized model of the ribosome exit tunnel and Sophie Bekisz for her help in drawing figure 1 (right panel). This work was supported by the FNRS-FWO EOS grant *n°*30480119 (Join-t-against-Osteoarthritis), the FNRS-WELBIO (THERAtRAME) in Belgium and the European Research Council under the European Union’s Horizon 2020 Framework Program (H2020/2014-2020) /ERC grant agreement *n°*772418 (INSITE).

## Notes

### Competing Interest Statement

The authors have declared no competing interest.

## References

[1] Marina Rodnina and Wolfgang Wintermeyer, “Fidelity of aminoacyl-tRNA selection on the ribosome: kinetic and structural mechanism.” Annu Rev Biochem 70, 415–35 (2001).

[2] Malte Beringer and Marina Rodnina, “The ribosomal peptidyl transferase,” Molecular Cell 26, 311–21 (2007).

[3] Poul Nissen, Jeffrey Hansen, Nenad Ban, Peter B. Moore, and Thomas A. Steitz, “The structural basis of ribosome activity in peptide bond synthesis,” Science 289, 920–930 (2000).

[4] Carolyn T. MacDonald, Julian H. Gibbs, and AC Pipkin, “Kinetics of biopolymerization on nucleic acid templates,” Biopolymers 6, 1–25 (1968).

[5] Carolyn T. MacDonald and Julian H. Gibbs, “Concerning the kinetics of polypeptide synthesis on polyribosomes,” Biopolymers 7, 707–725 (1969).

[6] Tobias von der Haar, “Mathematical and computational modelling of ribosomal movement and protein synthesis: an overview,” Computational and Structural Biotechnology Journal 1, e201204002 (2012).

[7] Hadas Zur and Tamir Tuller, “Predictive biophysical modeling and understanding of the dynamics of mRNA translation and its evolution,” Nucleic Acids Research 44(2016).

[8] Nicholas T. Ingolia, Liana F Lareau, and Jonathan S Weissman, “Ribosome profiling of mouse embryonic stem cells reveals the complexity and dynamics of mammalian proteomes,” Cell 147, 789–802 (2011).

[9] Tatsuya Morisaki, Kenneth Lyon, Keith F. DeLuca, Jennifer G. DeLuca, Brian P. English, Zhengjian Zhang, Luke D. Lavis, Jonathan B. Grimm, Sarada Viswanathan, Loren L. Looger, Timothee Lionnet, and Timothy J. Stasevich, “Real-time quantification of single RNA translation dynamics in living cells,” Science 352, 1425–1429 (2016).

[10] Aaron Fluitt, Elsje Pienaar, and Hendrik Viljoen, “Ribosome kinetics and aa-tRNA competition determine rate and fidelity of peptide synthesis,” Comput Biol Chem 31, 335–346 (2007).

[11] Jianli Lu and Carol Deutsch, “Electrostatics in the ribosomal tunnel modulate chain elongation rates,” Journal of Molecular Biology 384, 73–86 (2008).

[12] Tamir Tuller, Isana Veksler-Lublinsky, Nir Gazit, Martin Kupiec, Eytan Ruppin, and Michal Ziv-Ukelson, “Composite effects of gene determinants on the translation speed and density of ribosomes,” Genome Biology 12, R110 (2011).

[13] Marina Rodnina, “The ribosome in action: Tuning of translational efficiency and protein folding,” Protein Science 25(2016).

[14] Ajeet Sharma, Nabeel Ahmed, and Edward O’Brien, “Determinants of translation speed are randomly distributed across transcripts resulting in a universal scaling of protein synthesis times,” Phys Rev E 97(2018).

[15] Premal Shah, Yang Ding, Malwina Niemczyk, Grzegorz Kudla, and Joshua B. Plotkin, “Rate-limiting steps in yeast protein translation,” Cell 153, 1589–1601 (2013).

[16] Andrea Riba, Noemi Di Nanni, Nitish Mittal, Erik Arhné, Alexander Schmidt, and Mihaela Zavolan, “Protein synthesis rates and ribosome occupancies reveal determinants of translation elongation rates,” Proc Nat Acad Sci USA 116, 15023–15032 (2019).

[17] Alexandra Dana and Tamir Tuller, “The effect of tRNA levels on decoding times of mRNA codons,” Nucleic Acids Research 42(2014).

[18] Alon Raveh, Michael Margaliot, Eduardo D. Sontag, and Tamir Tuller, “A model for competition for ribosomes in the cell,” Journal of The Royal Society Interface 13, 20151062 (2016).

[19] Michael Y. Pavlov, Richard E. Watts, Zhongping Tan, Virginia W. Cornish, Måns Ehrenberg, and Anthony C. Forster, “Slow peptide bond formation by proline and other N-alkylamino acids in translation,” Proc Nat Acad Sci USA 106, 50–54 (2009).

[20] Khanh Dao Duc and Yun Song, “The impact of ribosomal interference, codon usage, and exit tunnel interactions on translation elongation rate variation,” PLOS Genetics 14, e1007166 (2018).

[21] Jian-Rong Yang, Xiaoshu Chen, and Jianzhi Zhang, “Codon-by-codon modulation of translational speed and accuracy via mRNA folding,” PLOS Biology 12, 1–14 (2014).

[22] Lisa J. Simpson, Ellie Tzima, and John S. Reader, “Mechanical forces and their effect on the ribosome and protein translation machinery,” Cells 9(2020).

[23] Thomas Gorochowski, Zoya Ignatova, Roel Bovenberg, and Hans Roubos, “Trade-offs between tRNA abundance and mRNA secondary structure support smoothing of translation elongation rate,” Nucleic Acids Research 43(2015).

[24] Carlo Artieri and Hunter Fraser, “Accounting for biases in riboprofiling data indicates a major role for proline in stalling translation,” Genome Research 24(2014).

[25] Renana Sabi and Tamir Tuller, “A comparative genomics study on the effect of individual amino acids on ribosome stalling,” BMC Genomics 16(2015).

[26] Rodrigo Requião, Henrique Souza, Silvana Rossetto, Tatiana Domitrovic, and Fernando Palhano, “Increased ribosome density associated to positively charged residues is evident in ribosome profiling experiments performed in the absence of translation inhibitors,” RNA Biology 13(2016).

[27] Catherine A. Charneski and Laurence D. Hurst, “Positively charged residues are the major determinants of ribosomal velocity,” PLOS Biology 11, 1–20 (2013).

[28] Neil Voss, Mark Gerstein, Thomas Steitz, and Peter Moore, “The geometry of the ribosomal polypeptide exit tunnel,” Journal of Molecular Biology 360, 893–906 (2006).

[29] Jianli Lu, William Kobertz, and Carol Deutsch, “Mapping the electrostatic potential within the ribosomal exit tunnel,” Journal of Molecular Biology 371, 1378–91 (2007).

[30] Carlos Pace, Gerald Grimsley, and J Martin Scholtz, “Protein ionizable groups: pK values and their contribution to protein stability and solubility,” The Journal of biological chemistry 284, 13285–9 (2009).

[31] Jin-Der Wen, Laura Lancaster, H. Courtney Hodges, Ana Zeri, Shige Yoshimura, Harry Noller, Carlos Bustamante, and Ignacio Tinoco, “Following translation by single ribosomes one codon at a time,” Nature 452, 598–603 (2008).

[32] Christian Kaiser and Ignacio Tinoco, “Probing the mechanisms of translation with force,” Chemical Reviews 114(2014).

[33] Tingting Liu, Ariel Kaplan, L. Alexander, Shannon Yan, Jin-Der Wen, L. Lancaster, C. E. Wickersham, K. Fredrick, H. Noller, I. Tinoco, and C. Bustamante, “Direct measurement of the mechanical work during translocation by the ribosome,” eLife 3(2014).

[34] Varsha Desai, Filipp Frank, Antony Lee, Maurizio Righini, Laura Lancaster, Harry Noller, Ignacio Tinoco, and Carlos Bustamante, “Co-temporal force and fluorescence measurements reveal a ribosomal gear shift mechanism of translation regulation by structured mRNAs,” Molecular Cell 75(2019).

[35] Leah Shaw, R Zia, and Kelvin Lee, “Totally asymmetric exclusion process with extended objects: A model for protein synthesis,” Phys Rev E, Statistical, nonlinear, and soft matter physics 68, 021910 (2003).

[36] R Zia, Jiajia Dong, and B Schmittmann, “Modeling translation in protein synthesis with TASEP: A tutorial and recent developments,” J Statist Phys 144(2011).

[37] Philip Greulich, Luca Ciandrini, Rosalind Allen, and Maria Romano, “Mixed population of competing totally asymmetric simple exclusion processes with a shared reservoir of particles,” Phys Rev E, Statistical, nonlinear, and soft matter physics 85, 011142 (2012).

[38] Khanh Dao Duc, Sanjit Batra, Nicholas Bhattacharya, Jamie Cate, and Yun Song, “Differences in the path to exit the ribosome across the three domains of life,” Nucleic Acids Research 47(2019).

[39] John David Jackson, Classical Electrodynamics (Wiley & Sons, (1998)) p. 32, third edition.

[40] Del Lucent, Christopher Snow, Colin Aitken, and Vijay Pande, “Non-bulk-like solvent behavior in the ribosome exit tunnel,” PLoS Computational Biology 6, e1000963 (2010).

[41] Harry Noller, L. Lancaster, J. Zhou, and J.P. Donohue, “The ribosome as an RNA-based molecular machine,” in EMBO Workshop:Protein Synthesis and Translational Control in EMBL Heidelberg, Germany (4-7 Sept 2019).

[42] Riccardo Belardinelli, Heena Sharma, Frank Peske, Wolfgang Wintermeyer, and Marina Rodnina, “Translocation as continuous movement through the ribosome,” RNA Biology 13, 1–7 (2016).

[43] Srividya Mohan, John Donohue, and Harry Noller, “Molecular mechanics of 30S subunit head rotation,” Proc Nat Acad Sci USA 111(2014).

[44] M.T. Friberg, P. Gonnet, Y. Barral, N.N. Schraudolph, and G.H. Gonnet, “Measures of codon bias in yeast, the tRNA pairing index and possible DNA repair mechanisms,” in P. Bucher and B. Moret (eds), Proceedings of the 6th Workshop on Algorithms in Bioinformatics (WABI), vol. 4175 of Lecture Notes in Bioinformatics (Springer Verlag, Berlin) (2006

[45] Gina Cannarozzi, Nicol Schraudolph, Mahamadou Faty, Peter von Rohr, Markus Friberg, Alexander Roth, Pedro Gonnet, Gaston Gonnet, and Yves Barral, “A role for codon order in translation dynamics,” Cell 141, 355–67 (2010).

[46] Carol Deutsch, “Tunnel vision: Insights from biochemical and biophysical studies,” in In: ItoK. (eds) Regulatory Nascent Polypeptides. Springer, Tokyo. (2014).

[47] Danny M. Hatters, “Protein misfolding inside cells: The case of huntingtin and Huntington’s disease,” IUBMB Life 60, 724–728 (2008).

[48] Montserrat Arrasate and Steven Finkbeiner, “Protein aggregates in Huntington’s disease,” Experimental neurology 238, 1–11 (2011).

[49] Mingchen Chen and Peter Wolynes, “Aggregation landscapes of huntingtin exon 1 protein fragments and the critical repeat length for the onset of Huntington’s disease,” Proc Nat Acad Sci USA 114, 201702237 (2017).

[50] Silvia Bonfanti, Maria Chiara Lionetti, Maria Fumagalli, Venkat Chirasani, Guido Tiana, Nikolay Dokholyan, Stefano Zapperi, and Caterina La Porta, “Molecular mechanisms of heterogeneous oligomerization of huntingtin proteins,” Scientific Reports 9(2019).

[51] Benjamin Fritch, Andrey Kosolapov, Phillip Hudson, Daniel A. Nissley, H. Lee Woodcock, Carol Deutsch, and Edward P. O’Brien, “Origins of the mechanochemical coupling of peptide bond formation to protein synthesis,” Journal of the American Chemical Society 140, 5077–5087 (2018).

[52] Daniel Goldman, Christian Kaiser, Anthony Milin, Maurizio Righini, Ignacio Tinoco, and Carlos Bustamante, “Mechanical force releases nascent chain-mediated ribosome arrest in vitro and in vivo,” Science 348, 457–60 (2015).

[53] Michael Thommen, Wolf Holtkamp, and Marina V Rodnina, “Co-translational protein folding: progress and methods,” Current Opinion in Structural Biology 42, 83–89 (2017).

[54] Rodrigo D. Requião, Luiza Fernandes, Henrique José Araujo de Souza, Silvana Rossetto, Tatiana Domitrovic, and Fernando L. Palhano, “Protein charge distribution in proteomes and its impact on translation,” PLOS Computational Biology 13, 1–21 (2017).

[55] Carlos Bustamante, Yann R. Chemla, Nancy R. Forde, and David Izhaky, “Mechanical processes in biochemistry,” Annual Review of Biochemistry 73, 705–748 (2004).

[56] Jordi Ribas-Arino and Dominik Marx, “Covalent mechanochemistry: Theoretical concepts and computational tools with applications to molecular nanomechanics,” Chemical Reviews 112, 5412–5487 (2012).

[57] GI Bell, “Models for the specific adhesion of cells to cells,” Science 200, 618–627 (1978).

[58] Marina Rodnina, Andreas Savelsbergh, Vladimir Katunin, and Wolfgang Wintermeyer, “Hydrolysis of GTP by elongation factor G drives tRNA movement on the ribosome,” Nature 385, 37–41 (1997).

