## Supplementary figures and images for "Ribosome exit tunnel electrostatics"

### Supplemental Figure 1: animated figure of ribosome exit tunnel

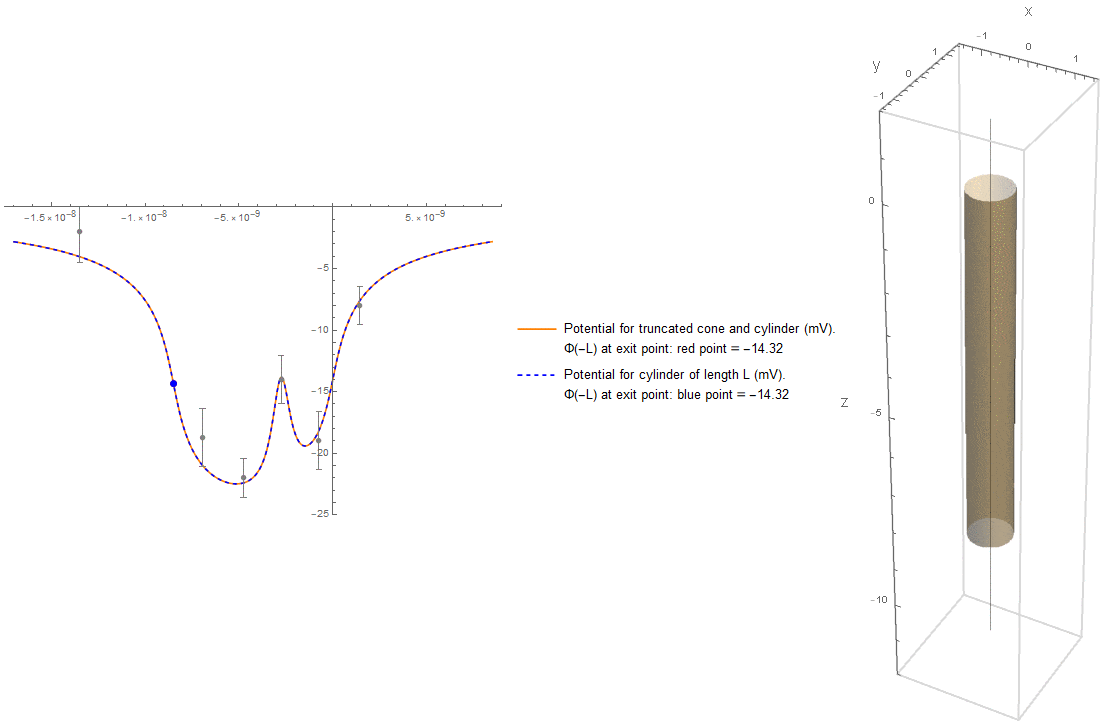
